# A Geometric Optical Analysis of Human Retinal Cells and the Hypothesis of Monocular Stereoscopic Vision

**DOI:** 10.1101/2025.01.02.631067

**Authors:** Yong Ding, Dalu Ding, Dawei Ding, Rongzhi Jiang

**Affiliations:** Agape Medical Technologies Co., Ltd.; Jmuse Technologies Co., Ltd.

## Abstract

We propose a geometric optical model that describes real-image formation via monocular vision in three dimensions. We only assume that the ganglion cell nuclei, bipolar cell nuclei, and photoreceptor cell nuclei in the human retina can be treated as microlenses and actively participate in imaging. In particular, we suggest that monocular vision operates via an array of high-magnification micro-real-image telescopes oriented in various azimuthal directions. The main elements of each telescope are the following. The first is the cornea and crystalline lens, which serve as objective lenses with tunable focal lengths. The second is a paired array of ganglion cell nuclei and bipolar cell nuclei, which act as a conjugate mirror. The third is an individual photoreceptor cell nuclei (cone or rod nuclei), which function as a micro-eyepiece. The fourth is the outer segment of each photoreceptor cell, which serves as a micro-object-distance sensor.

We also hypothesize that six combinations of conjugate mirrors and micro-eyepiece arrays work within the human eye. Each combination is responsible for perceiving different ranges of object distances. We further hypothesize that the process by which one human eye perceives object distances involves two stages: from near to finite distances, and from finite distances to infinite distances. In different stages, the distance between different optical elements is tuned so that the object distance can be computed a specific formula when the photocurrent at the retina photoreceptor’s outer segment layer vanishes.

Lastly, we conduct computational simulations. We find that there are two possible processes. By tuning the distances between the elements of our optical model, the eye can perceive object distances as close as 25 cm to dozens of meters in one process and dozens of meters to kilometers in the other process, depending on the sensitivity of the eye to small changes in focal plane spacing.

## 1. Introduction

When Charles Wheatstone invented the stereoscope in 1838, it has been believed that human stereoscopic vision primarily comes from both eyes. This is achieved by perceiving differences in the positions (or angles) of images of objects at various distances on the retinas of the two eyes [1,2]. For monocular stereoscopic vision, it is generally thought that the brain estimates depth by comparing the sizes of known objects on the retina.

We also already know [3] that the human eye can see objects through the process in which the cornea and crystalline lens, acting as an objective lens, form an image of light from external objects onto the retina. The photoreceptor cells on the retina respond to light by generating a photocurrent, and the brain’s optic nerve processes the photocurrent information to achieve vision. Our current understanding is that a single photoreceptor cell only identifies a single bright spot on the two-dimensional retina where the image is formed.

This paper proposes that the above understanding of monocular stereoscopic vision and the optical functions of the cells in the retina (such as ganglion cells, bipolar cells, photoreceptor cells, and others) is still insufficient. To this day, our understanding of the human eye and brain remains very limited.

In this article, by assuming that the ganglion cell nuclei, bipolar cell nuclei, and photoreceptor cell nuclei (cone cell nuclei and rod cell nuclei) on the retina function as optical microlenses and also participate in optical imaging, a geometric optical model for three-dimensional real-image formation in monocular human vision is proposed. This model suggests that our single eye functions as an array of high-magnification miniature real-image telescopes oriented toward numerous azimuthal angles (each azimuthal angle has two angular coordinates). This miniature real-image telescope array is composed of the following fundamental optical components: the cornea and crystalline lens serving as the objective lens, ganglion cell nuclei arrays and bipolar cell nuclei arrays forming phase conjugate mirrors, the photoreceptor cell nuclei arrays forming micro-eyepiece arrays, and the photoreceptor outer segment array forming a micro-distance sensor array. The two angular coordinates of the azimuthal angle of each miniature real-image telescope’s micro-eyepiece or micro-distance sensor correspond to the two angular coordinates of the azimuthal angle of a three-dimensional object in space. Each micro-distance sensor is a photoreceptor outer segment of multiple photosensitive layers at varying depths, where each depth corresponds to the object’s distance in space. This article also speculates on the process of monocular object distance perception in the human eye. Using this three-dimensional real-image optical model, a formula for perceived object distance values is derived, and numerical simulations of object distance perception are conducted for distances ranging from 25 cm to finite distances, as well as from finite distances to infinite distances.

## 2. Results

This section concerns the monocular three-dimensional optical imaging model, hypothesis of the stereoscopic vision perception process, and derivation of the formula for object distance perception.

### 2.1. Monocular Three-Dimensional Optical Imaging Model This paper assumes

1. The ganglion cell nuclei, bipolar cell nuclei, and photoreceptor cell nuclei (cone cell nuclei and rod cell nuclei) on the retina are also optical elements, functioning as optical microlenses (referred to as “cell lenses”). Their optical properties can be approximated as ideal thin microlenses with two coincident principal planes. The layer composed of a large number of cell nuclei forms a cell lens array.

Based on the above basic assumption and referencing the anatomical structure of the human retina [4,5], Figure 1 (Figure 1A and Figure 1B) illustrates the monocular three-dimensional imaging optical model of the human eye proposed in this paper. This schematic diagram represents a certain optical meridional plane (or a specific elevation angle in a three-dimensional spherical coordinate system, θ= 0). The fundamental optical components are:

**Figure 1:**
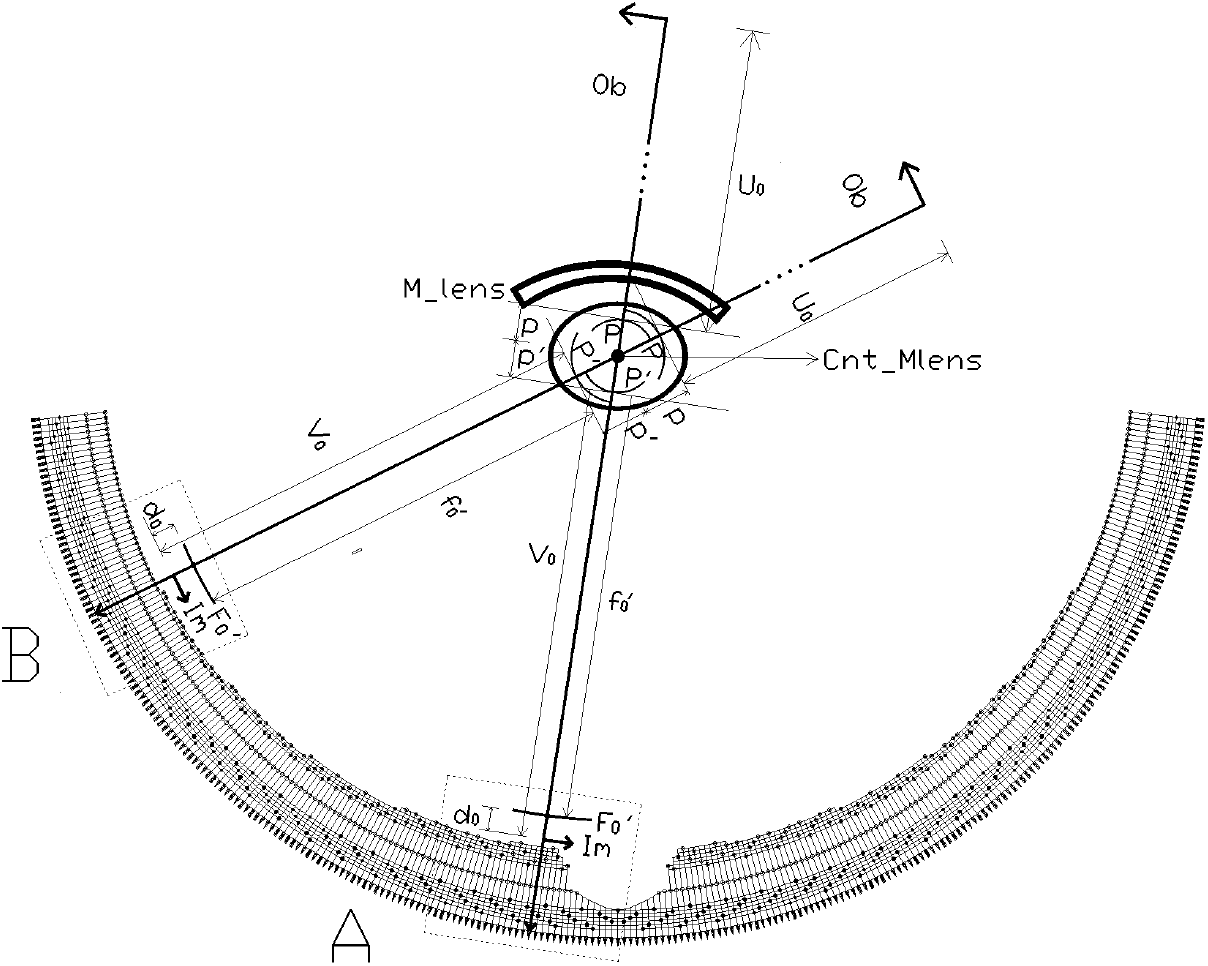
Schematic diagram of the three-dimensional optical imaging model of a single eye. [**M_lens**: Objective lens composed of cornea + crystalline lens; **Cnt_Mlens**: Center of the objective lens; ***P, P***′: Principal planes of the objective lens on the object side and the image side; ***p, p***′: Distance from the center of the objective lens to the principal planes on the object side and the image side; **Ob**: Object in three-dimensional space; ***U***_**0**_, ***V***_**0**_: Object distance and image distance of the objective lens; 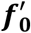 : Focal length on the image side of the objective lens; 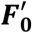 : Focal plane on the image side of the objective lens; **Im**: First image; ***d***_**0**_: Distance between the image and the focal plane on the image side of the objective lens]

**Figure 1A:**
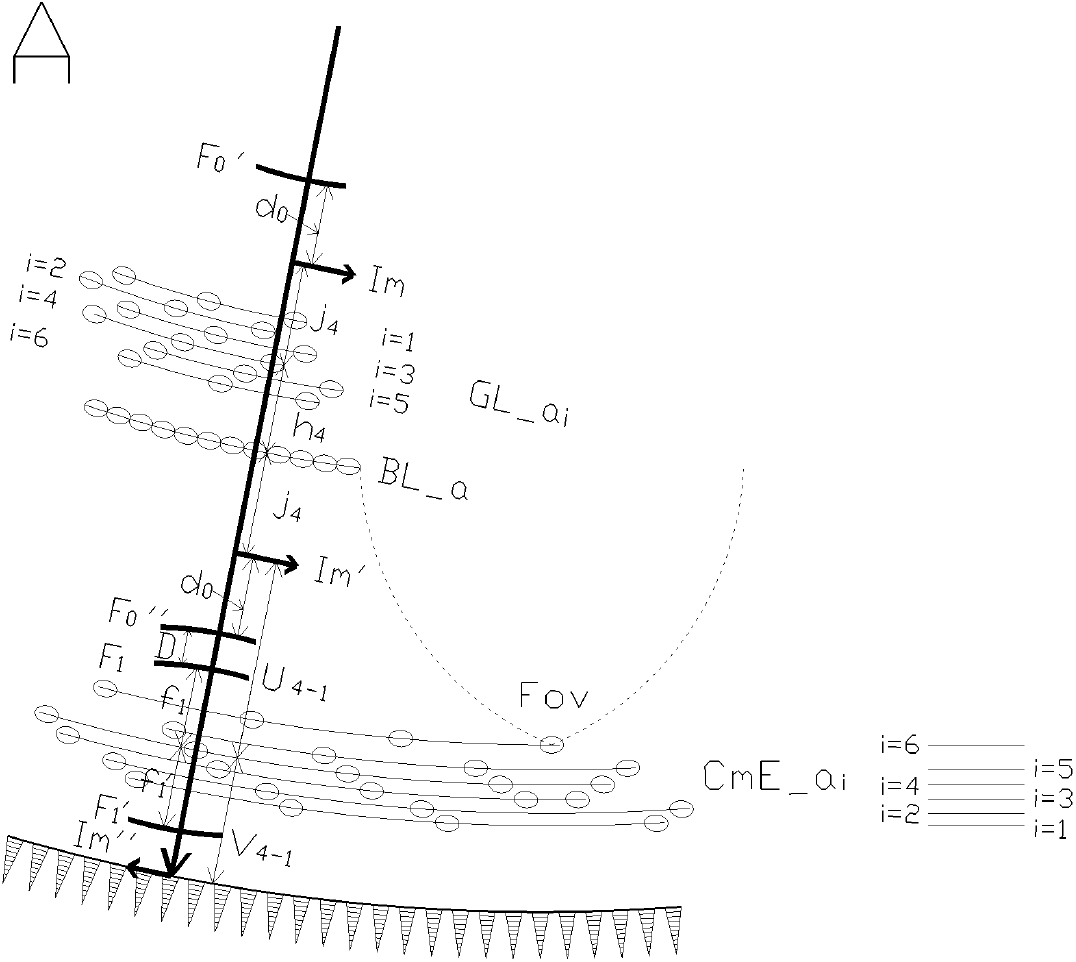
Schematic diagram of the three-dimensional optical imaging model at the fovea centralis. [**i**: Index; **Fov**: Fovea centralis; ***F*0**’: Image-side focal plane of the objective lens; **Im**: The first image (always coincides with a conjugate plane. In this figure, it coincides with the conjugate plane of conjugate mirror 4); ***d***_**0**_: Distance between the image and the image-side focal plane of the objective lens; **GL_ai (i=1**,**2**,…,**6)**: Six ganglion cell lens arrays; **BL_a**: Bipolar cell lens array; **hi (i=1**,**2**,…,**6)**: interlayer spacings between the six ganglion cell arrays and the single bipolar cell lens array (only ***h***_**4**_ is labeled in the figure); ***j***_***i***_ **(i=1**,**2**,…,**6)**: Conjugate distances of the six conjugate mirrors (only ***j***_**4**_ is labeled in the figure); **Im’**: Conjugate image of **Im**; 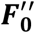: Image-side conjugate focal plane of the objective lens after passing through conjugate mirror 4; ***F***_**1**_, ***F***_**1**′_ : Object-side and image-side focal planes of the cone-cell micro-eyepiece; ***f***_**1**_, ***f***_**1**_′: Object-side and image-side focal lengths of the cone-cell micro-eyepiece; ***D***: Distance between 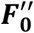: and ***F***_**1**_; ***U***_***i***−**1**_, ***V***_***i***−**1**_ **(i=1**,**2**,…,**6)**: Object distance and image distance of the cone lens at the first photosensitive layer (only **i=4** is labeled in the figure); **Im”**: Image of **Im’** after passing through the cone-cell micro-eyepiece; **CmE_ai (i=1**,**2**,…,**6)**: Six cone-cell micro-eyepiece arrays]

**Figure 1B:**
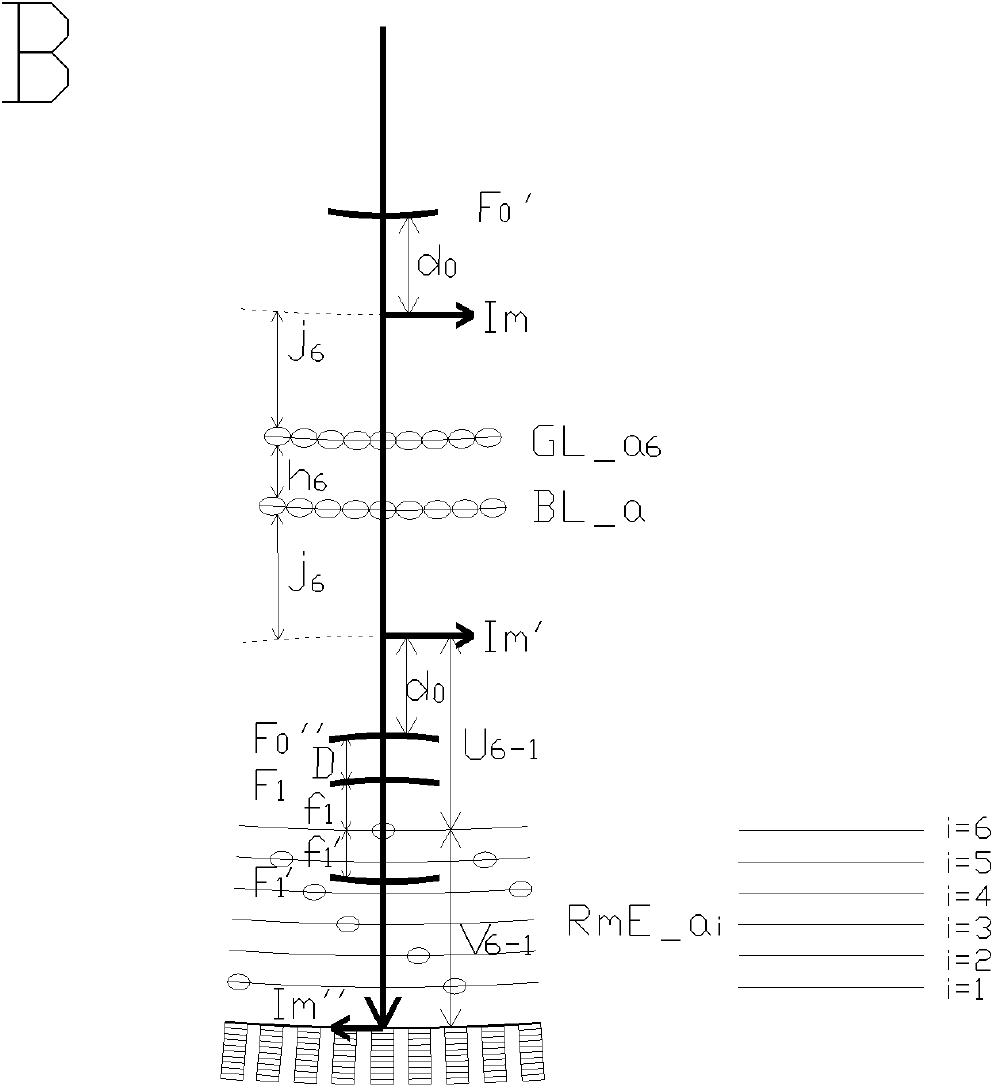
Schematic diagram of the three-dimensional optical imaging model at a location away from the fovea centralis. [**i**: Index; Image-side focal plane of the objective lens; **Im**: The first image (always coincides with a conjugate plane. In this figure, it coincides with the conjugate plane of conjugate mirror 6); **d0**: Distance between the image and the focal plane; **GL_a6**: Ganglion cell lens array 6; **BL_a**: Bipolar cell lens array; ***h*_6_**: interlayer spacing between ganglion cell lens array 6 and the bipolar cell lens array; ***j*_6_**: Conjugate distance of conjugate mirror 6; **Im’**: Conjugate image of **Im**; ***F*_0_′′** : Image-side conjugate focal plane of the objective lens after passing through conjugate mirror 6; ***F*_1_, *F*_1_′** : Object-side and image-side focal planes of the rod-cell micro-eyepiece; ***f*_1_**, ***f*_1_′** : Object-side and image-side focal lengths of the rod-cell micro-eyepiece; ***D***: Distance between ***F*0**′′ and ***F*_1_**; ***U*_i=1_**, ***V_i−1_* (i=1**,**2**,…,**6)**: Object distance and image distance of the rod lens at the first photosensitive layer (only i=6 is labeled in the figure); **Im”**: Image of **Im’** after passing through the rod-cell micro-eyepiece; **RmE_ai (i=1,2**,…,**6)**: Six rod-cell micro-eyepiece arrays]

1. Objective Lens: The cornea and crystalline lens together form the objective lens. In this model, the focal plane and principal plane of the objective lens are spherical. Its focal length is continuously adjustable within a certain range.
2. Conjugate Mirrors: A large number of ganglion cell nuclei and bipolar cell nuclei, distributed spherically with the center of the objective lens as the sphere’s center, form six ganglion cell lens arrays and one bipolar cell lens array, respectively. Each of the six ganglion cell lens arrays pairs with the bipolar cell lens array to form six conjugate mirrors (Figure 1,Figure 1A). Along a single radial line (in terms of azimuth) on the image side (inside the human eye) with the center of the objective lens as the origin, there is only one ganglion cell lens from one of the six layers of ganglion cell lens arrays, combined with one bipolar cell lens. It is assumed that, from the fovea to a position 10 cm away from the fovea, the number of ganglion cell lens arrays decreases from six to one, at a rate of one array fewer for every 2 mm. Consequently, the number of conjugate mirrors also decreases sequentially. At a distance of 10 mm or farther from the fovea, there is only one conjugate mirror remaining (Figure 1,Figure 1B).
3. Micro-Eyepieces: A photoreceptor cell nucleus forms a photoreceptor lens, which acts as a micro-eyepiece. Along a single radial line (in terms of azimuth) with the center of the objective lens as the origin, there is only one micro-eyepiece. The azimuths of all micro-eyepieces in space do not overlap. The azimuth of a micro-eyepiece corresponds one-to-one with the object angle in three-dimensional space. A cone cell nucleus forms a cone-cell micro-eyepiece, while a rod cell nucleus forms a rod-cell micro-eyepiece. A large number of photoreceptor cell nuclei, distributed spherically with the objective lens as the center, form six micro-eyepiece arrays. It is assumed that from the fovea to a position 10 mm away from the fovea, the number of rod-cell micro-eyepieces increases from 0% to 100% of the total at a rate of a 20% increase every 2 mm. At all positions, the cone-cell micro-eyepieces and rod-cell micro-eyepieces are evenly distributed across the six micro-eyepiece arrays. Near the fovea, all micro-eyepieces in the six micro-eyepiece arrays are composed of cone cell nuclei and are referred to as cone-cell micro-eyepiece arrays. At a distance of 10 mm or farther from the fovea, all micro-eyepieces in the six micro-eyepiece arrays are composed of rod cell nuclei and are referred to as rod-cell micro-eyepiece arrays.
4. Micro-Object Distance Sensor: A single photoreceptor outer segment with multiple layers of photosensitive membranes at different depths constitutes a micro-object distance sensor. A three-dimensional real image is formed on the photosensitive membranes at different depths within the micro-object distance sensor. The photosensitive membrane at a specific depth detects the light from the image and generates a photocurrent. The depth of the photosensitive membrane that generates the photocurrent indirectly represents the distance of the external object in three-dimensional space. The farther a photosensitive membrane is from the micro-eyepiece, the greater the corresponding object distance. The multiple layers of photosensitive membranes in a single micro-object distance sensor are arranged in parallel. Each micro-object distance sensor is combined with one micro-eyepiece from the 6 micro-eyepiece arrays, and this pair lies on a straight line or shares a specific azimuthal angle with respect to the optical center of the objective lens. The azimuthal angles of all micro-object distance sensors are spatially non-overlapping. The azimuthal angle of a micro-object distance sensor corresponds one-to-one with the angular position of an object in three-dimensional space. A large number of photoreceptor outer segments, distributed spherically around the optical center of the objective lens, form an array of micro-object distance sensors. Near the fovea, the majority of photoreceptor cells are cone cells, and the micro-object distance sensor array is composed of cone-based micro-object distance sensors. At locations 10 mm or farther from the fovea, the majority of photoreceptor cells are rod cells, and the micro-object distance sensor array is composed of rod-based micro-object distance sensors. Assuming that the proportion of cone cells decreases from 100% at the fovea to 0% at 10 mm away from the fovea, in which the proportion decreases by 20% for every 2 mm, the proportion of cone-based micro-object distance sensors in the array also changes correspondingly. A single objective lens, a conjugate mirror, and a micro-eyepiece combined with a micro-object distance sensor along the same azimuth (defined by two angular coordinates) together form a micro-real-image telescope. A micro-real-image telescope can form a three-dimensional real image of a single object point within its azimuth or of objects within a very small angular range, and it can perceive the distance of the object(s). An array of micro-real-image telescopes, distributed across different azimuths and centered on the optical center of the objective lens (forming a spherical arrangement), enables the perception of object distances in three-dimensional space.

### 2.2. Hypothesis on the Process of Monocular Stereoscopic Perception

To form a hypothesis on how distance perception is achieved with a single eye, the following assumptions are made based on the three-dimensional imaging model illustrated in Figures 1,1A, and 1B:

1. A single micro-real-image telescope perceives the distance of only one object point within a specific azimuth (defined by two angular coordinates).
2. The object-side conjugate plane of the conjugate mirror always coincides with the image plane of the first image. The first image, after passing through the conjugate mirror, forms the conjugate image of the first image.
3. The conjugate mirror pairs with six micro-eyepiece arrays to form six conjugate micro-eyepiece array combinations. 3-A) Near the fovea, six conjugate mirrors with different conjugate distances are combined one-to-one with six cone-cell micro-eyepiece arrays to form six conjugate cone-cell micro-eyepiece array combinations (denoted as JJ-*C*_*i*_, where i = 1, 2, …, 6 is the combination index). For example: The conjugate mirror with the shortest conjugate distance (conjugate mirror 1) pairs with the outermost cone-cell micro-eyepiece array (micro-eyepiece array 1), which is responsible for perceiving the range of the nearest object distances, forming conjugate cone-cell micro-eyepiece array combination 1 (JJ-*C*_6_). The conjugate mirror with the longest conjugate distance (conjugate mirror 6) pairs with the innermost cone-cell micro-eyepiece array (micro-eyepiece array 6), which is responsible for perceiving the range of the farthest object distances, forming conjugate cone-cell micro-eyepiece array combination 6 (JJ-*C*_6_). 3-B) In the retinal region away from the fovea, a conjugate mirror with the longest conjugate distance (conjugate mirror 6) pairs with six rod-cell micro-eyepiece arrays to form six conjugate rod-cell micro-eyepiece array combinations (denoted as JJ-*R*_*i*_, where i = 1, 2, …, 6, is the combination index). For example: conjugate mirror 6 pairs with the outermost rod-cell micro-eyepiece array (micro-eyepiece array 1), which is responsible for perceiving the range of the nearest object distances, forming conjugate rod-cell micro-eyepiece array combination 1 (JJ-*R*_1_). conjugate mirror 6 pairs with the innermost rod-cell micro-eyepiece array (micro-eyepiece array 6), which is responsible for perceiving the range of the farthest object distances, forming conjugate rod-cell micro-eyepiece array combination 6 (JJ-*R*_6_).
4. All ganglion cell lenses, bipolar cell lenses, cone-cell lenses, and rod-cell lenses have the same focal length within their respective types. However, the focal lengths of these different types of lenses may either differ from one another or be the same.
5. For each micro-eyepiece array, the image distance (the distance from the micro-eyepiece center to the first photosensitive layer) when the photocurrent disappears, denoted as *V*_i− 1_, is fixed. Similarly, the corresponding object distance (the distance from the conjugate image to the micro-eyepiece center), denoted as *U*_i−1_, is also fixed. (Alternatively, this implies that the arrangement of photoreceptor cells and the distance between the nuclei of photoreceptor cells and their outer segments remains constant.)
6. The total distance *L* from the center of the objective lens to the first photosensitive layer of the micro-object distance sensor (the layer closest to the nuclei of the photosensitive cells) is a fixed constant for a given individual. Alternatively, their relative positions are fixed and unchanging.
7. Define the focal plane spacing *D* as the distance between the image-side focal plane of the objective lens after passing through the conjugate mirror, and the object-side focal plane of the micro-eyepiece. When *D* = 0, the conjugate mirror as a whole (e.g., in the case of the fovea, the six conjugate mirrors act as a whole; in regions away from the fovea, a single conjugate mirror acts as a whole) can move up or down relative to the micro-eyepiece array. The movement distance is represented by the change in q, which is the distance from the bipolar cell lens array of the conjugate mirror to the first photosensitive layer. Each conjugate micro-eyepiece combination has its own fixed q.
8. The distance *p* (or *p*′) from the center of the objective lens to the principal plane changes with the focal length of the objective lens. The maximum and minimum values of *p*′ (where minimum *p*′ = 0) constrain the perceivable object distance range for each conjugate micro-eyepiece array combination when *D* = 0 and *L* is fixed.
9. When the focal plane spacing *D* > 0, changes in *D* are achieved through the synchronized changes in *h*_+_ (the distance between the ganglion cell array and the bipolar cell array of conjugate mirror 6) and *q*.

Based on the above assumptions, this article hypothesizes that the perception of object distance is realized through two processes: From near to finite distances, and from finite distance to infinite distances.

1. The process of perceiving object distance from near to finite distances: This is achieved by maintaining the condition where the conjugate focal plane of each objective lens in each conjugate micro-eyepiece array combination coincides with the focal plane of the corresponding micro-eyepiece (*D* = 0), while scanning and adjusting the focal length of the objective lens.

The specific steps for perceiving object distance from near to finite distance can be completed using one of the following three methods:

1-A) Starting with conjugate cone-cell micro-eyepiece array combination 1 or conjugate rod-cell micro-eyepiece array combination 1, the focal length of the objective lens is scanned from small to large values. When the focal length of the objective lens in a certain combination reaches a specific value, the human brain perceives that one or more micro-distance sensors at certain azimuthal angles generate photocurrent. The focal length of the objective lens at this moment is termed the “photocurrent generation focal length.” At this point, the real image in the micro-real-image telescope is located deep within the micro-object distance sensor, and the human brain perceives the presence of an object.

As the focal length of the objective lens continues to increase to a certain focal length, the photocurrent disappears. The focal length of the objective lens at this moment is termed the “photocurrent disappearance focal length.” At this point, the real image in the micro-real-image telescope is located on the first photosensitive layer of the micro-object distance sensor. The human brain (or nervous system) determines the object distance based on:

- Which conjugate mirror and micro-eyepiece array combination is being used, and
- The value of the photocurrent disappearance focal length.

The brain then records the three-dimensional visual information of the perceived object distance, including:

1. The two angular coordinates of the azimuthal position of the micro-object distance sensor that perceived the object, and
2. The object distance itself.

1-B) The human brain, based on experience, estimates the object distance. Based on the estimated object distance, it selects a certain conjugate micro-eyepiece array combination. The focal length of the objective lens is scanned from small to large values until, at a certain focal length of the objective lens in a certain combination, the human brain perceives that one or more micro-distance sensors at certain azimuthal angles generate photocurrent. At this moment, the real image in the micro-real-image telescope is located deep within the micro-distance sensor, and the human brain perceives the presence of an object. When the focal length of the objective lens continues to increase to a certain focal length, the photocurrent disappears. At this moment, the real image in the micro-real-image telescope is located on the first photosensitive layer of the micro-distance sensor. The human brain calculates the object distance based on which conjugate mirror and micro-eyepiece array combination is used and the value of the photocurrent disappearance focal length and records the three-dimensional visual information of the perceived object distance, including the two angular coordinates of the azimuthal position of the micro-distance sensor that perceived the object and the object distance itself.

1-C) After the human eye locates the target object through the above process 1-A or 1-B, the focal length of the objective lens stays near the photocurrent disappearance focal length. The focal length is finely adjusted through multiple back-and-forth scans to obtain the precise value of the photocurrent disappearance focal length. The human brain calculates the precise object distance based on which conjugate mirror and micro-eyepiece array combination is used and the precise value of the photocurrent disappearance focal length and records the three-dimensional visual information of the perceived precise object distance, including the two angular coordinates of the azimuthal position of the micro-distance sensor that perceived the object and the precise object distance itself.

For the above process of perceiving object distance from near to finite distance, based on *D* = 0 and scanning and adjusting the focal length of the objective lens, the six conjugate cone-cell micro-eyepiece array combinations and the six pairs of conjugate rod-cell micro-eyepiece array combinations each have their own object distance perception range.

2)The process of perceiving object distance from finite distance to infinity: This is achieved by fixing the focal length of the objective lens and scanning to increase the distance between the conjugate focal plane of the objective lens and the focal plane of the micro-eyepiece, i.e., focal plane spacing *D* > 0. Specifically:

2-1) Using the conjugate micro-eyepiece array combination 6 (either the conjugate cone-cell micro-eyepiece array combination 6 or the conjugate rod-cell micro-eyepiece array combination 6) with the longest object distance perception capability, the focal length of the objective lens is set to its maximum value. The focal length of the objective lens is then fixed, and the total distance *L* is also kept fixed. This corresponds to the maximum object distance perceivable by the conjugate micro-eyepiece array combination 6 when *D* = 0. This maximum value serves as the starting value for the process of perceiving object distance from finite distance to infinity.

2-2) By scanning to increase the distance between the conjugate focal plane of the objective lens and the focal plane of the micro-eyepiece, i.e., increasing *D* from *D* = 0, the perceivable object distance range is extended. This scanning to increase *D* is achieved by the human brain scanning to compress the interlayer spacing *h*_+_ between the ganglion cell layer and the bipolar cell layer that constitute conjugate mirror 6, and performing a scanning operation to move conjugate mirror 6 as a whole away relative to the photoreceptor cell nuclear layer. The distance scanned and moved is represented by the increase in *q*, which is the distance from the bipolar cell lens array of the conjugate mirror to the first photosensitive layer.

The specific steps for perceiving object distance from finite distance to infinity can be described using the increase of the focal plane spacing *D*: *D* starts from *D* = 0 and increases incrementally until, at a certain *D*, the human brain perceives that one or more micro-distance sensors at certain azimuthal angles generate photocurrent. This value of *D* is named the “photocurrent generation focal plane spacing.” At this moment, the real image in the micro-real-image telescope is located deep within the micro-distance sensor, and the human brain perceives the presence of an object. When D continues to increase to a certain value, the photocurrent disappears. This value of *D* is named the “photocurrent disappearance focal plane spacing.” At this moment, the real image in the micro-real-image telescope is located on the first photosensitive layer of the micro-distance sensor. The human brain (nervous system) calculates the object distance based on which conjugate mirror and micro-eyepiece array combination is used and the “photocurrent disappearance focal plane spacing,” and records the three-dimensional visual information of the perceived object distance, including the two angular coordinates of the azimuthal position of the micro-distance sensor that perceived the object and the object distance itself.

2-3) When *D, h*_+_, and *q* are simultaneously adjusted to approach certain (maximum or minimum) extreme values, specifically near *D*_∞_, *h*6min, and *q*max, the perceivable object distance increases dramatically. This results in the images of all objects beyond a certain distance being focused on the first photosensitive layer. At this point, the photocurrent begins to fluctuate intermittently. The human brain perceives these intermittent fluctuations in the photocurrent to recognize that there are objects beyond a certain distance. However, it is unable to determine the exact distance of these objects or distinguish the distance differences between objects at this farthest range. This specific distance is referred to as the “farthest perceivable object distance for monocular vision.” This farthest distance is related to the brain’s ability (or resolution) to precisely distinguish the values of *D, h*6, and *q*.

The human brain (nervous system) calculates the object distance based on which conjugate mirror and micro-eyepiece array combination is used and the focal plane spacing at which the photocurrent fluctuates intermittently. It records the three-dimensional visual information of the perceived object distance, including the two angular coordinates of the azimuthal position of the micro-distance sensor that perceived the object and the object distance itself.

The conjugate cone-cell micro-eyepiece array combination 6 and the conjugate rod-cell micro-eyepiece array combination 6 have different starting values for perceiving object distance, as well as different values for *D*_∞_, *h*_6min_, and *q*_max_.

This article further hypothesizes that the human brain undergoes the following learning process to determine the constants required for perceiving object distances from near to finite distances and from finite to infinite distances. These constants include which conjugate micro-eyepiece combination to use, the *q* value for each combination, the focal length of the objective lens, the adjustment range of the focal plane spacing, and so on:

1. The brain selects one conjugate micro-eyepiece array combination at a time and scans the entire range of object distances by either increasing the focal length of the objective lens from small to large or scanning to increase the focal plane spacing *D*. This process is used to preliminarily determine these constants.
2. The above process is repeated multiple times to precisely determine these constants.

## 3. Results

### 3.1. Derivation and Simulation of the Formula for Monocular Perception of Object Distance

In the following formula, the meanings of the various parameter symbols are described in the caption of Figure 1. The distance from the object-side principal plane to the center of the objective lens, the object distance and the object-side focal length of the objective lens, and the object distance and the object-side focal length of the cellular lens, are all represented as negative values [2].

Based on the hypothesized process of object distance perception described in Section 2.2, the derivation of the formula for the human brain’s perception/calculation of object distance is as follows:

1. Using a certain micro-real-image telescope, as its objective lens focal length or focal plane spacing is tuned from small to large, when a specific focal length or focal plane spacing is reached, the human brain perceives the generation of a photocurrent: it perceives the presence of an object. As shown in Figures 1,1A,and 1B, at this moment, an object within the range of a small object angle (defined by two object angle coordinates) forms an inverted image (the first image) through the objective lens along the optical path of the micro-real-image telescope, located within the vitreous body near the retina. A conjugate mirror is then adjusted so that its object-side conjugate plane coincides with the position of the first image plane. At this point, the first image is flipped (Figure 1 shows a vertical flip) and magnified by the conjugate mirror (Considering the significant difference—hundreds of times— between the spacing of the two cellular layers that make up the conjugate mirror and the focal length of the objective lens, we can assume that these two cellular layers are parallel up to a small difference in angle. Therefore, this paper assumes that the conjugate mirror does not have a magnification function.). This is the image within the retina formed by the conjugate mirror, which is still an inverted conjugate image. This inverted conjugate image is located farther along the object-side focal point of the micro-eyepiece. The inverted conjugate image in turn via the micro-eyepiece forms another real image, which is an upright image of the three-dimensional object. When the photocurrent is just generated, the real image is located deeply in the micro-photosensitive layer of the micro-object-distance sensor (farther than the photoreceptor cell nuclei).
2. When the focal length or focal plane spacing of the objective lens continues to increase to a certain focal length or focal plane spacing, the position of the real image on the micro-object-distance sensor moves from the deeper layer to the first photosensitive layer. As this focal length or distance increases further, the photocurrent begins to disappear. When this happens, the focal length or focal plane spacing of the objective lens is referred to as the photocurrent disappearance focal length or photocurrent disappearance focal plane spacing. At this moment, the object distance *U*_0_ of the objective lens can be calculated using the formula (1) derived from formulas (1a) to (1f).

The imaging formula of the objective lens:

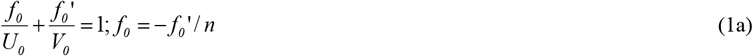

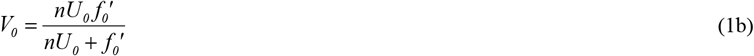

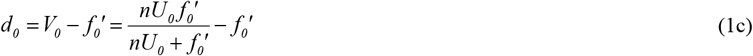

The imaging formula of the micro-eyepiece:

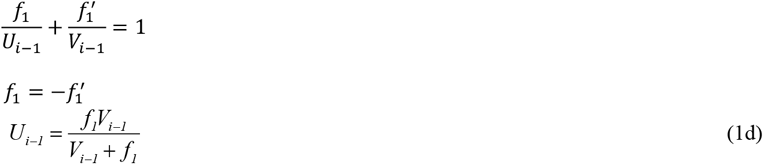

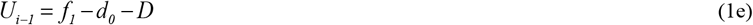

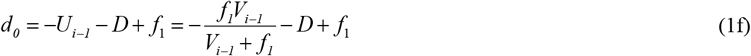

Combining the above formula, we obtain

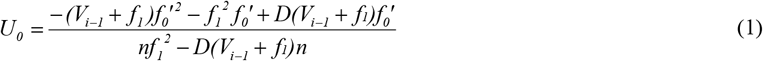

For the conjugate mirror *i*, composed of the ganglion cell lens array *i* and the bipolar cell lens array as shown in Figures 1A and 1B, the relationship between the interlayer spacing of the two cellular lens arrays, *h*_*i*_, and their conjugate distance,*ji* (both *h*_*i*_ and*j*_*i*_ are defined as positive values), is as follows:

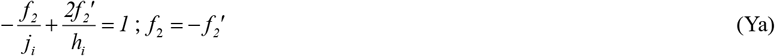

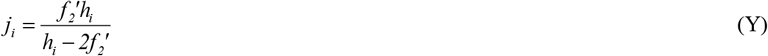

Here, *f2*_2_ and 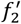 represent the object-side and image-side focal lengths of the cellular lenses, respectively (assuming that the focal lengths of the ganglion cell lenses and bipolar cell lenses are equal).

Let the total distance from the center of the objective lens to the first photosensitive layer of the object-distance sensor be *L*, and let the distance from the center of the objective lens to the image-side principal plane be p’. This can be calculated using formula (2):

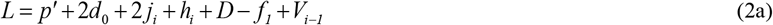

Substitute equation (1f) for *d*_0_ into equation (2a) to obtain

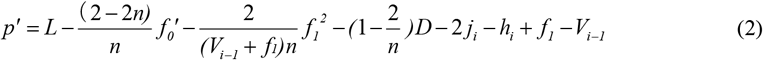

The distance q from the bipolar cell lens array of the conjugate mirror to the first photosensitive layer can be calculated using formula (3):

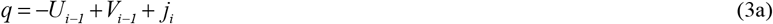

Substitute equation (1d) for *U*_*i* −1_ into equation (3a) to obtain

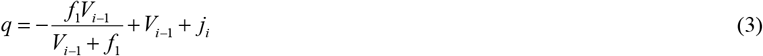

Equations (1), (2), and (3) indicate that once *V*_*i* −1_, *f*_1_, *n, D, L, h*, and *j*_*i*_ are fixed, for each conjugate micro-eyepiece array combination *i*:

1. *U*_0_ is uniquely determined by 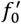,
2. *p*′ is also uniquely determined by 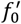,
3. *q*, however, is independent of both 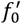 and *D*, and is instead a constant related only to the specific conjugate micro-eyepiece array combination being used (which depends on *V*_*i* −1_ and *j*_*i*_).

Table 1 provides estimated values of the geometric optical parameters for the lenses in the three-dimensional imaging optical model of a single eye. These include the objective lens (cornea and crystalline lens), ganglion cells, bipolar cells, cone cells, and rod cells [1,2]. Here, the estimated sizes and focal lengths of the ganglion cell lenses, bipolar cell lenses, and cone-cell lenses are assumed to be identical, while the sizes and focal length of the rod cell nucleus are half that of the other cellular lenses.

**Table 1:** Estimated geometric optical parameters of lenses in the three-dimensional imaging optical model of a single eye.

| Lens | Radius ($\mu\text{m}$) | Lens Refractive Index ($n_l$) | Environment Refractive Index ($n$) | Object-Side Focal Length ($f$) ($\mu\text{m}$) | Image-Side Focal Length ($f'$) ($\mu\text{m}$) |
| --- | --- | --- | --- | --- | --- |
| Objective Lens<br>(Cornea + Crystalline Lens) | - | - | Object side:<br>$n_0 = 1$<br>Image side: $n = 1.33$ | -11955 ~ -18496 | 15900 ~ 24600 |
| Ganglion Cell Lens | 2.535 | 1.42 | 1.33 | -20 | 20 |
| Bipolar Cell Lens | - | - | - | - | - |
| Cone-cell lens | - | - | - | - | - |
| Rod-cell lens | 1.268 | 1.42 | 1.33 | -10 | 10 |

Tables 2 and 3 list the following estimated data based on anatomical measurements of the human retina from the literature [4,5]:

**Table 2:**
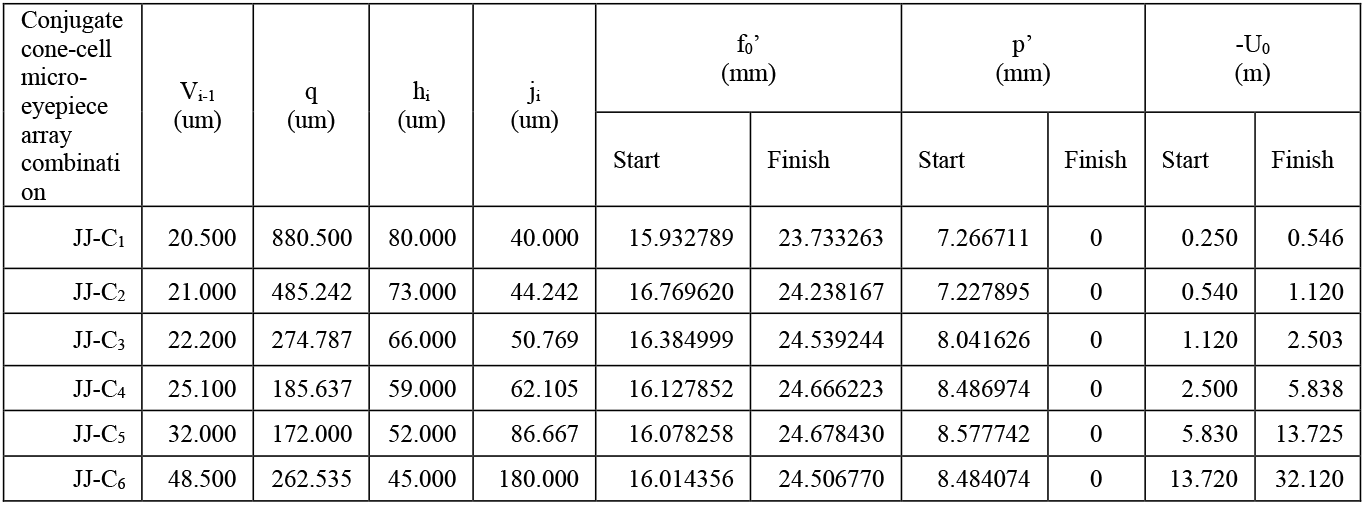
Key optical parameters of the six conjugate cone-cell micro-eyepiece array combinations for perception of object distances from near to finite distances, along with focal length adjustment and synchronization of the distance p′ (from the objective lens center to the image-side principal plane) with the range of perceived object distances.

**Table 3:** Key optical parameters of the six conjugate rod-cell micro-eyepiece array combinations for perception of object distances from near to finite distances, along with focal length adjustment and synchronization of the distance p′ (from the objective lens center to the imageside principal plane) with the range of perceived object distances.

| Conjugate rod-cell micro-eyepiece array combination | $V_{i-1}$<br>($\mu\text{m}$) | $q$<br>($\mu\text{m}$) | $h_i$<br>($\mu\text{m}$) | $j_i$<br>($\mu\text{m}$) | $f_0'$<br>(mm) | | $p'$<br>(mm) | | $-U_0$<br>(m) | |
| --- | --- | --- | --- | --- | --- | --- | --- | --- | --- | --- |
|  |  |  |  |  | Start | Finish | Start | Finish | Start | Finish |
| JJ-R<sub>1</sub> | 20.500 | 220.024 | 45 | 180 | 16.009102 | 24.551782 | 8.536350 | 0 | 20.200 | 47.500 |
| JJ-R<sub>2</sub> | 26.000 | 222.250 |  |  | 19.889949 | 24.546470 | 4.656551 | 0 | 47.500 | 72.340 |
| JJ-R<sub>3</sub> | 32.000 | 226.545 |  |  | 20.933693 | 24.543929 | 3.610217 | 0 | 72.340 | 99.440 |
| JJ-R<sub>4</sub> | 38.000 | 231.571 |  |  | 21.756096 | 24.540536 | 2.783761 | 0 | 99.440 | 126.520 |
| JJ-R<sub>5</sub> | 44.000 | 236.941 |  |  | 22.270324 | 24.535146 | 2.264794 | 0 | 126.520 | 153.560 |
| JJ-R<sub>6</sub> | 50.000 | 242.500 |  |  | 22.620392 | 24.530013 | 1.909608 | 0 | 153.560 | 180.580 |

1. The distances *V*_*i*−1_ (where *i* = 1, 2, 3, …, 6) from the centers of the cone and rod cell micro-eyepiece arrays to the first photosensitive layer of the micro-object-distance sensor. Note that the estimated values of *V*_*i*−1_ differ for the cone and rod cell arrays.
2. The six interlayer spacings *h*_*i*_ (where *i* = 1, 2, 3, …, 6) between the six ganglion cell lens arrays and the single bipolar cell lens array.
3. The conjugate distances *j*_*i*_ for the six conjugate mirrors, calculated using the optical parameter estimates from Table 1 and Equation (Y) [6–9].

### 3.2A. Calculation and simulation of monocular perception of distances from near to finite distances

During the process of perceiving distances from near to finite distances, it is speculated in this paper that the conjugate focal plane position of the objective lens coincides with the object-side focal plane of the eyepiece, with a focal plane spacing *D* = 0; the distance from the center of the objective lens to the first photosensitive layer, *L*, is fixed. At this point, equations (1) and (2) are rewritten as:

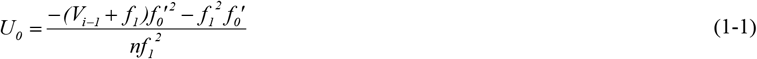

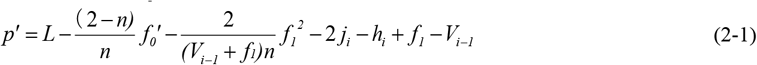

Formula (1-1) indicates that when *D* = 0, there is a parabolic relationship between *U*_0_ and 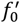. For each conjugate rod-cell micro-eyepiece array combination *i*, the further the distance *U*_0_, the longer 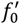 is needed for perception. Formula (2-1) shows that *p*′ decreases as 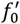 increases. In the case of a finite and fixed L, the 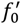 when *p*′ = 0 and its corresponding *U*_0_ represent the maximum focal length of the objective lens and the furthest perceivable distance for that specific conjugate micro-eyepiece array combination.

Figures 2,3,and 4 represent the relationships between the objective lens’s photocurrent disappearance focal length 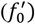, the distance from the objective lens center to the image-side principal point/plane (*p*′), the distance from the bipolar cell lens array of the conjugate mirror to the first photosensitive layer (*q*), and the perceived distance *U*_0_ for six conjugate cone-cell micro-eyepiece array combinations. These figures are based on the estimation of geometric optical parameters of lenses using formulas (1-1), (2-1), (3), and the values from tables 1 and 2,with *D* = 0 and *L* = 25 cm. Table 2 also shows the value of *q* and adjustable ranges for 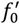 and *p*′ and their corresponding ranges for the perceivable distances *U*_0_.

**Figure 2:**
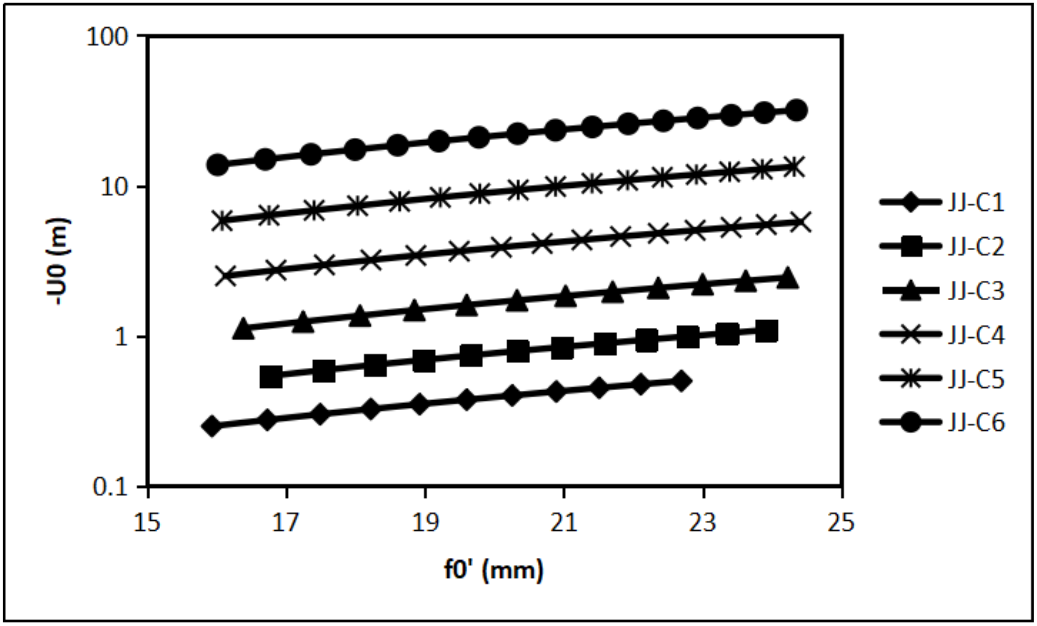
Using six conjugate cone-cell micro-eyepiece array combinations, along with the optical parameter estimations of the lenses in Table 1,the conjugate mirror parameters in Table 2,and under the conditions *D* = 0 and *L* = 25 cm, the relationship between the photocurrent disappearance focal length of the objective lens 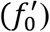 and the perceived object distance (*U*_0_) is demonstrated. In the figure, the *U*_0_ axis is presented on a logarithmic scale in meters.

**Figure 3:**
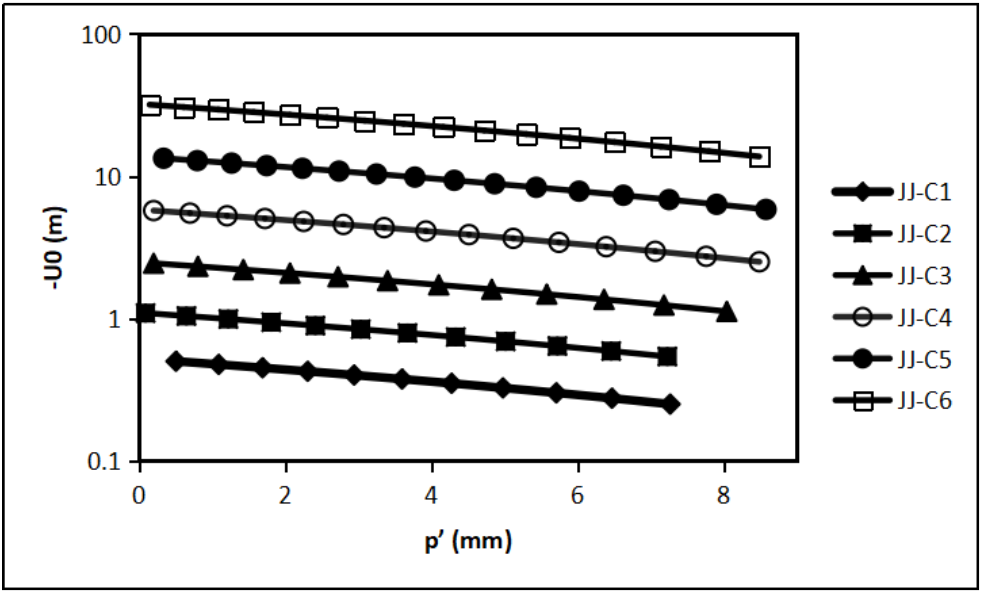
Using six conjugate cone-cell micro-eyepiece array combinations, along with the optical parameter estimations of the lenses in Table 1,the conjugate mirror parameters in Table 2,and under the conditions *D* = 0 and *L* = 25 cm, the relationship between the distance from the center of the objective lens to the image-side optical principal point/plane (*p*′) and the perceived object distance (*U*_0_) is demonstrated. In the figure, the *U*_0_ axis is presented on a logarithmic scale in meters.

**Figure 4:**
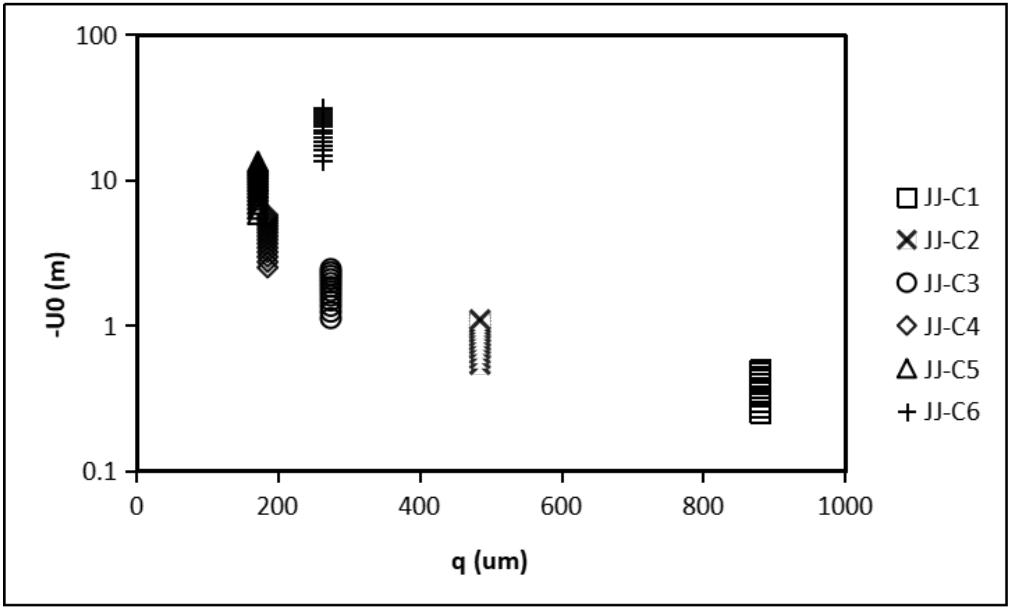
Using six conjugate cone-cell micro-eyepiece array combinations, along with the estimates of the lenses’ optical parameters in Table 1,the conjugate mirror parameters in Table 2,and under the conditions *D* = 0 and *L* = 25 cm, the relationship between the distance from the bipolar cell lens array of the conjugate mirror to the first photosensitive layer (*q*) and the perceived object distance (*U*_0_) is demonstrated. In the figure, the *U*_0_ axis is presented on a logarithmic scale in meters.

Figure 2 and Table 2 indicate that for each conjugate cone-cell micro-eyepiece array combination, the objective lens’s photocurrent disappearance focal length, 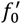, needs to increase as the perceivable object distance increases. The outermost layer of the cone-cell micro-eyepiece arrays (farthest from the objective lens center) surrounding the fovea, specifically cone-cell micro-eyepiece array 1 combined with conjugate mirror 1 (conjugate cone-cell micro-eyepiece array combination 1), can perceive objects as close as 0.25 m. The innermost cone-cell micro-eyepiece array 6 combined with conjugate mirror 6 (conjugate cone-cell micro-eyepiece array combination 6) can perceive objects as far as 32.12 m. The focal length of the objective lens can range approximately from 15.9 to 24.7 mm.

Figure 3 and Table 2 show that for each conjugate cone-cell micro-eyepiece array combination, the distance from the center of the objective lens to the image-side optical principal point/plane, *p*′, needs to decrease as the perceivable object distance increases. At the maximum perceivable object distance for each combination, *p*′ reduces to 0 (the focal length of the objective lens is the maximum value for that combination). At the minimum perceivable object distance for each combination, *p*′ reaches its maximum value (the focal length of the objective lens is the minimum value for that combination), which is approximately 7.2–8.6 mm.

Figure 4 and Table 2 indicate that, for each conjugate cone-cell micro-eyepiece array combination, the distance *q* from the bipolar cell lens array of the conjugate mirror to the first photosensitive layer has a fixed value. When using conjugate cone-cell micro-eyepiece array combination 1, which perceives the smallest object distance, *q* is at its maximum: approximately 880.5 µm. When using conjugate cone-cell micro-eyepiece array combination 5, *q* is at its minimum: approximately 172 µm. This result also suggests that, to perceive different object distance ranges, the conjugate mirror as a whole needs to be moved to the corresponding position based on the specific conjugate cone-cell micro-eyepiece array combination being used. The range of movement required for the conjugate mirror is approximately 880.5 - 172 = 708.5 µm.

Figures 5,6,and 7 represent the relationships between the objective lens’s disappearance focal length 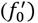, the distance from the objective lens center to the image-side principal point/plane (*p*′), the distance from the bipolar cell lens array of the conjugate mirror to the first photosensitive layer (*q*), and the perceived distance *U*_0_ for six conjugate rod-cell micro-eyepiece array combinations. These figures are based on the estimation of geometric optical parameters of lenses using formulas (1-1), (2-1), (3), and the values from Tables 1 and 3,with *D* = 0 and *L* = 25 cm.

**Figure 5:**
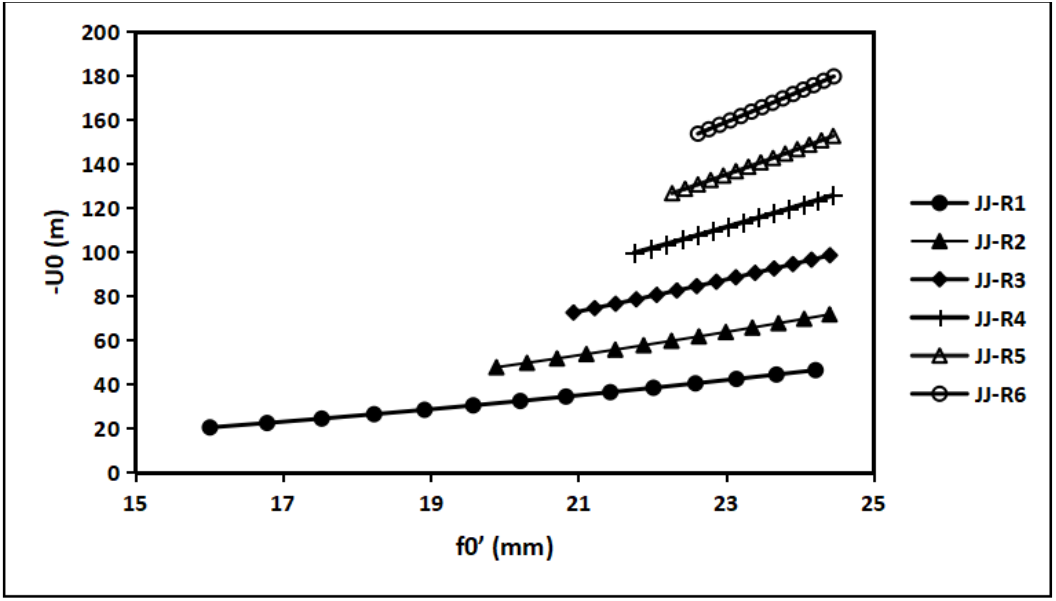
Using six conjugate rod-cell micro-eyepiece array combinations, along with the optical parameter estimations of the lenses in Table 1,the conjugate mirror parameters in Table 3,and under the conditions *D* = 0 and *L* = 25 cm, the relationship between the objective lens’s disappearance focal length 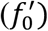 and the perceived object distance (*U*_0_) is demonstrated.

**Figure 6:**
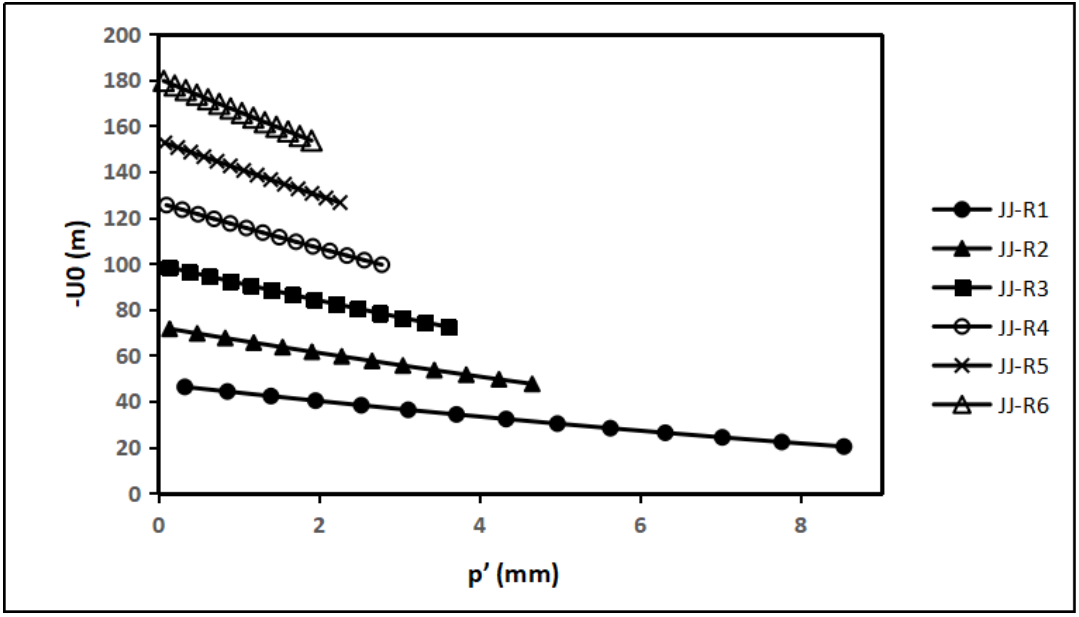
Using six conjugate rod-cell micro-eyepiece array combinations, along with the optical parameter estimations of the lenses in Table 1,the conjugate mirror parameters in Table 3,and under the conditions *D* = 0 and *L* = 25 cm, the relationship between the distance from the center of the objective lens to the image-side optical principal point/plane (*p*′) and the perceived object distance (*U*_0_) is demonstrated.

**Figure 7:**
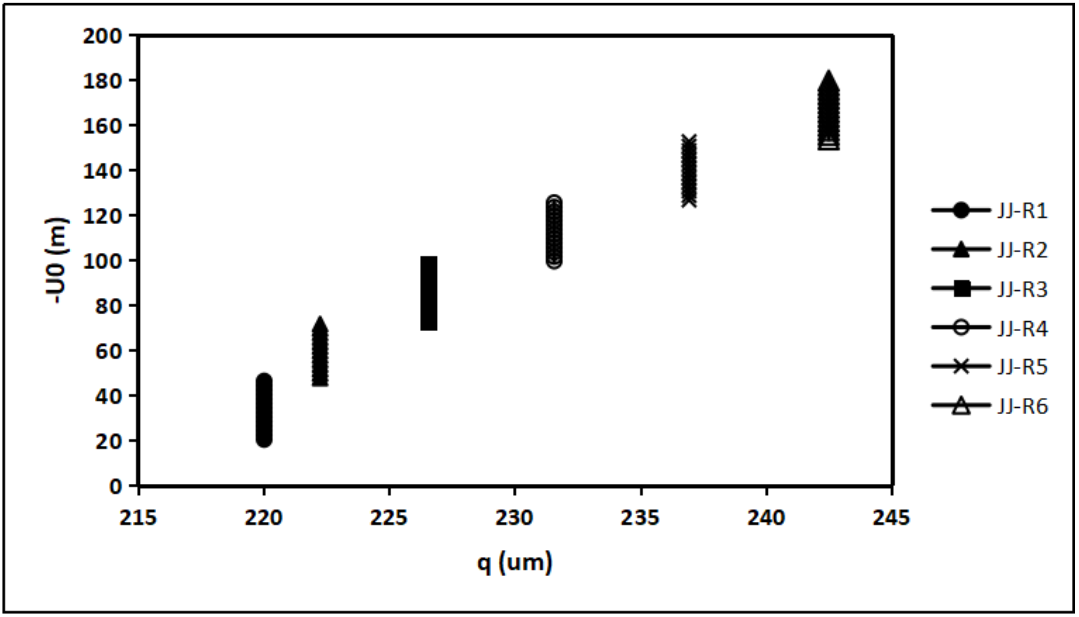
Using six conjugate rod-cell micro-eyepiece array combinations, along with the optical parameter estimations of the lenses in Table 1,the conjugate mirror parameters in Table 3,and under the conditions *D* = 0 and *L* = 25 cm, the relationship between the distance from the bipolar cell lens array of the conjugate mirror to the first photosensitive layer (*q*) and the perceived object distance (*U*_0_) is demonstrated.

Table 3 also shows the value of *q* and adjustable ranges for 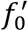 and *p*′ and their corresponding ranges for the perceivable distances *U*_0_.

Figure 5 and Table 3 indicate that for each conjugate rod-cell micro-eyepiece array combination, the objective lens’s photocurrent disappearance focal length, 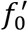, needs to increase as the perceivable object distance increases. The outermost (farthest from the objective lens center) rod-cell micro-eyepiece array 1 combined with conjugate mirror 6 (conjugate rod-cell micro-eyepiece array combination 1) can perceive objects at distances as close as 20.2 m. The innermost rod-cell micro-eyepiece array 6 combined with conjugate mirror 6 (conjugate rod-cell micro-eyepiece array combination 6) can perceive objects at distances as far as 180.58 m. The focal length of the objective lens can range approximately from 16.0 to 24.6 mm.

Figure 6 and Table 3 show that for each conjugate rod-cell micro-eyepiece array combination, the distance from the center of the objective lens to the image-side optical principal point/plane, *p*′, needs to decrease as the perceivable object distance increases. At the maximum perceivable object distance for each combination, *p*′ reduces to 0 (the focal length of the objective lens is the maximum value for that combination). At the minimum perceivable object distance for each combination, *p*′ reaches its maximum value (the focal length of the objective lens is the minimum value for that combination), which is approximately 1.9–8.5 mm.

Figure 7 and Table 3 indicate that, for each conjugate rod-cell micro-eyepiece array combination, the distance from the bipolar cell lens array of the conjugate mirror to the first photosensitive layer, *q*, has a fixed value. When using conjugate rod-cell micro-eyepiece array combination 1, which perceives the smallest object distance, *q* is at its maximum: approximately 220.0 µm. When using conjugate rod-cell micro-eyepiece array combination 5, *q* is at its minimum: approximately 242.5 µm. This result also suggests that, to perceive different object distance ranges, the conjugate mirror as a whole needs to be moved to the corresponding position based on the specific conjugate rod-cell micro-eyepiece array combination being used. The range of movement required for the conjugate mirror is approximately 242.5 - 220.0 = 22.5 µm.

### 3.2B. Simulation of monocular perception of finite to infinite object distances

Using conjugate micro-eyepiece array combination 6 (*i* = 6), when the distance between the conjugate focal plane of the objective lens and the object-side focal plane of the micro-eyepiece array, the focal plane spacing, is *D* = 0, and the focal length of the objective lens 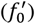 is adjusted to its maximum value 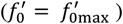, referred to as the “maximum objective lens focal length”, at this point (*p*′ = 0), and the perceivable object distance reaches its finite maximum limit, *U*_0max_ (referred to as the “maximum object distance”). If this 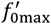 is fixed, and the total distance L is also fixed while increasing the focal plane spacing *D*, the relationship between the perceivable object distance *U*_0_ and *D* changes, and is modified from Equation (1) to:

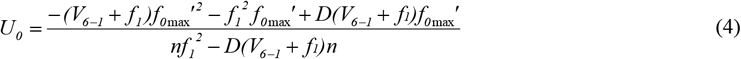

Equation (4) can be rewritten as

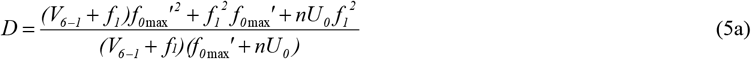

When *U*_0_ is infinitely large, from Equation (5a) we can obtain the corresponding focal plane spacing *D*_∞_:

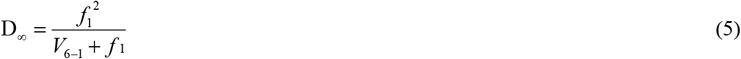

Using Equation (5), the focal length (*f*1) values of the cone-cell lens and the rod-cell lens from Table 1,as well as the *V*_6−1_ values (from Table 4), the maximum focal plane spacing (*D*_∞_) required for perceiving infinitely distant objects can be calculated. For the cone-cell lens, the maximum focal plane spacing is *D*_∞_ = 14.035 µm, and for the rod-cell lens, the maximum focal plane spacing is *D*_∞_ = 2.5000 μm.

**Table 4:** The fixed image distance of the cell lens, the fixed maximum focal length of the objective lens, the focal plane spacing, and other parameter ranges when using conjugate micro-eyepiece array combination 6 to perceive finite to infinite object distances.

| Conjugate micro-eyepiece Array Combination | $V_{6-1}$ (um) | $f'_{0max}$ (mm) | $D$ (um) | | $h_i$ (um) | | $j_i$ (um) | | $q$ (um) | | $-U_0$ (m) | |
| --- | --- | --- | --- | --- | --- | --- | --- | --- | --- | --- | --- | --- |
| | | | Initial $U_0$ | $\infty U_0$ | Initial $U_0$ | $\infty U_0$ | Initial $U_0$ | $\infty U_0$ | Initial $U_0$ | $\infty U_0$ | Initial $U_0$ ($U_{0max}$) | $\infty U_0$ |
| JJ-$C_6$ | 48.500 | 24.506770 | 0 | 14.035 | 45.000 | 44.786 | 180.000 | 187.124 | 194.464 | 201.589 | 32.120 | ∞ |
| JJ-$R_6$ | 50.000 | 24.530013 | 0 | 2.500 | 45.000 | 44.960 | 180.000 | 181.270 | 217.500 | 218.769 | 180.580 | ∞ |

Figures 8 and 9 respectively illustrate the relationship between the focal plane spacing (*D*) and the perceivable object distance (*U*_0_) when using the conjugate cone and rod-cell micro-eyepiece array combinations 6 (JJ-*C*_6_ and JJ-*R*_6_), corresponding to *V*_6−1_, with their 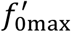 values fixed (as shown in Table 4). As *D* increases, the perceivable object distance (*U*_0_) rises sharply/exponentially, until it reaches infinity at their respective *D*_∞_.

**Figure 8:**
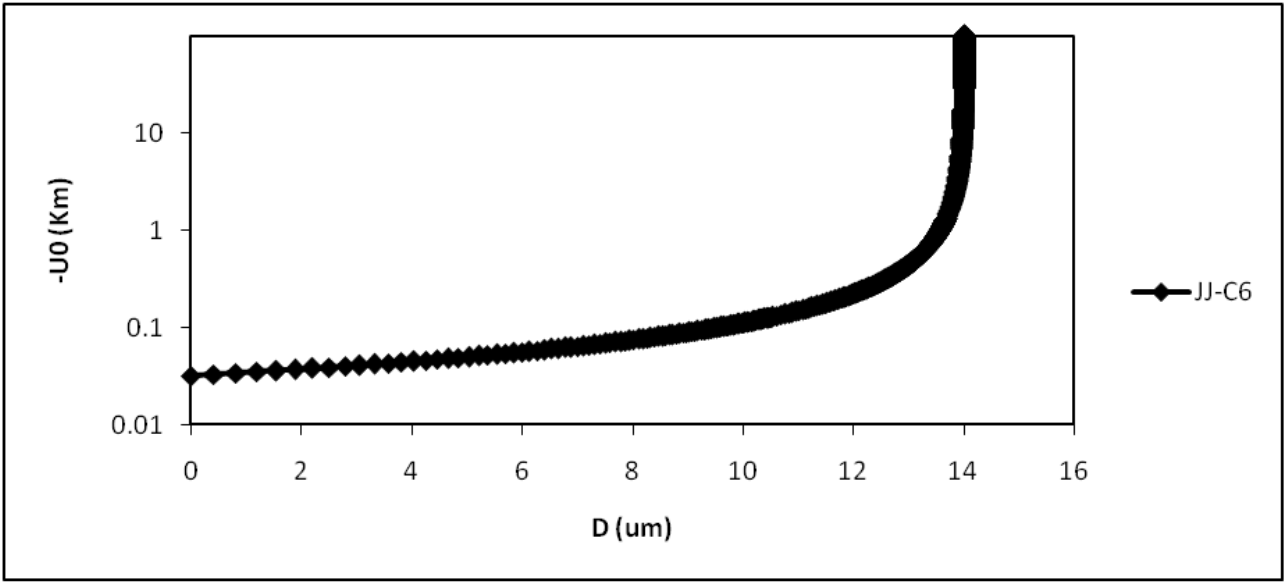
Using the conjugate cone-cell micro-eyepiece array combination 6, JJ-*C*_6_, corresponding to *V*_6−1_ and the 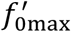 value (Table 4), the relationship between the focal plane spacing *D* and the perceivable object distance *U*_0_ (where the *U*_0_-axis is expressed in kilometers, log scale) is calculated based on Equation (4).

**Figure 9:**
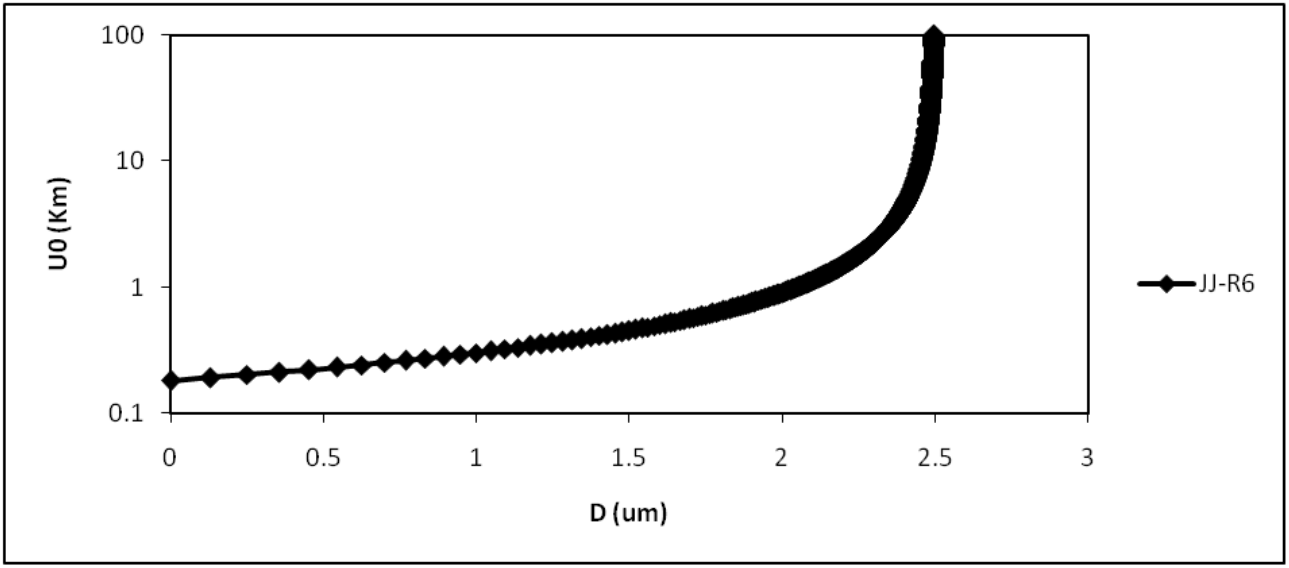
Using the conjugate rod-cell micro-eyepiece array combination 6, JJ_0_*R*_6_, corresponding to *V*_6−1_ and the 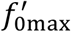 value (Table 4), the relationship between the focal plane spacing *D* and the perceivable object distance *U*_0_ (where the *U*_0_-axis is expressed in kilometers, log scale) is calculated based on Equation (4).

Let Δ*D* represent the difference between the focal plane spacing *D* and *D*,:

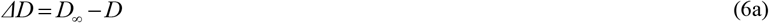

Substituting *D* from Equation (5a) and *D*_∞_ from Equation (5) into Equation (6a), we obtain:

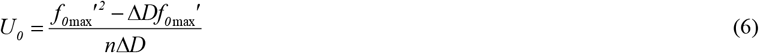

Equation (6) shows that if parameter Δ*D* is used for object distance perception, the perceivable object distance *U*_0_ is independent of the focal length *f*_1_ of the (cone or rod) cell lens. However, it should be noted that *D*_∞_, and *D* in Equation (6a), which define Δ*D*, are related to *f*_1_ (as shown in Equations (5) and (5a)).

Figure 10 and Table 5 illustrate the relationship between the perceivable object distance *U*_0_ and the difference Δ*D* from the maximum focal plane spacing when using the conjugate cone or rod-cell micro-eyepiece array combination 6, JJ-*C*_6_ or JJ-*R*_6_, with 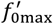 both set to 24.5 mm, as calculated using Equation (6). Figure 10 and Table 5 demonstrate that when using the conjugate micro-eyepiece array combination 6, the perceivable farthest object distance for a single eye decreases sharply from infinity as the focal plane spacing approaches the maximum focal plane spacing *D*_∞_, up to a few nanometers. As shown in Table 6, when the difference from *D*_∞_, is less than Δ*D* = 1, 10, or 20 nm, the perceivable farthest object distance reduces to 450, 45, and 23 km, respectively. This indicates that if the human brain can distinguish a difference of 1, 10, or 20 nm between *D* and *D*_∞_, it can perceive objects’ distances if the distance is less than 450, 45, and 23 km, respectively, instead of an infinite distance.

**Table 5:** Perceivable object distance when using conjugate micro-eyepiece array combination 6 and the difference from the maximum focal plane spacing is Δ*D* = 0 to 23 nm.

| $\Delta D(\text{nm})$ | $-U_0(\text{km})$ | $\Delta D$ (nm) | $-U_0(\text{km})$ | $\Delta D$ (nm) | $-U_0$ (km) | $\Delta D$ (nm) | $-U_0$ (km) |
| --- | --- | --- | --- | --- | --- | --- | --- |
| 0 | $+\infty$ | 6 | 75 | 12 | 38 | 18 | 25 |
| 1 | 450 | 7 | 64 | 13 | 35 | 19 | 24 |
| 2 | 225 | 8 | 56 | 14 | 32 | 20 | 23 |
| 3 | 150 | 9 | 50 | 15 | 30 | 21 | 21 |
| 4 | 113 | 10 | 45 | 16 | 28 | 22 | 20 |
| 5 | 90 | 11 | 41 | 17 | 26 | 23 | 20 |

**Figure 10:**
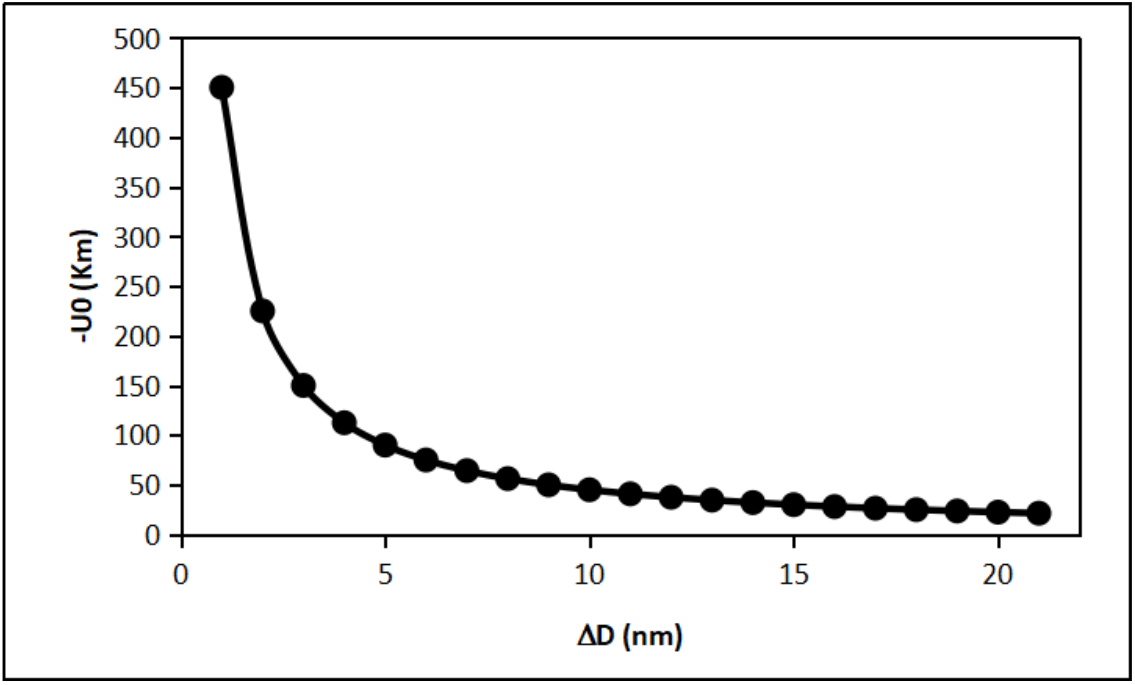
Using the conjugate cone or rod-cell micro-eyepiece array combination 6, JJ_0_C6 or JJ_0_ R6, with 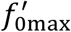 both being 24.5 mm, the relationship between the perceivable object distance *U*_0_ (represented in kilometers, log scale, on the *U*_0_-axis) and the difference Δ*D* of the maximum focal plane spacing is calculated based on Equation (6).

During the process of perceiving object distances from finite to infinite values, using the conjugate cone or rod-cell micro-eyepiece array combination 6 (JJ-*C*_6_ or JJ-*R*_6_), the focal length of the objective lens, 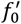, is fixed at its maximum value, 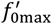 . At this point, the distance from the image-side conjugate focal plane of the objective lens to the conjugate image 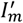, denoted as *L*, is also fixed.

At the maximum focal length of the objective lens:

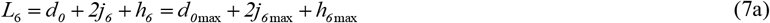

Here, 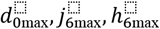 respectively represent *d*_0_, *j*_+_, *h*_+_ at the maximum focal length of the objective lens. Substituting *d*_0_ from Equation (1c) and *j*_+_ from Equation (Y) into Equation (7a), we obtain:

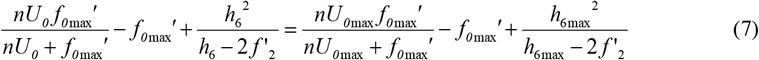

At this point, substituting *j*_+_ from Equation (Y) into Equation (3), the distance *q* from the bipolar cell lens array of the conjugate mirror to the first photosensitive layer is given as:

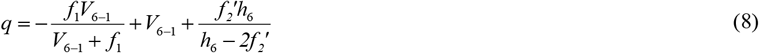

When *U*_0_ is at infinity, *h*_6_ reaches its minimum value, 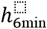, we have the following equation from equation (7):

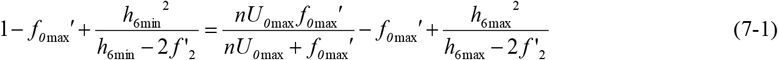

When *U*_0_ is at infinity, *h*_+_ reaches its minimum value, and *q* reaches its maximum value 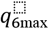, then we have the following equation from equation (8):

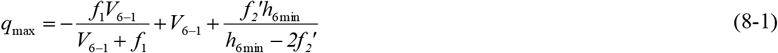

Table 4 shows the range of changes in *D, h*_6_, *j*_6_ and *q* during the process of perceiving object distances from finite to infinite values, calculated using Equations (7), (8), (7-1), and (8-1) for conjugate cone-cell micro-eyepiece array combination 6 and conjugate rod-cell micro-eyepiece array combination 6.

Figures 11 and 12 illustrate the relationships between *U*_0_, *h*_+_, and *q* during the process of perceiving object distances from finite to infinite values, calculated using Equations (7) and (8), respectively, when using conjugate cone-cell micro-eyepiece array combination 6.

**Figure 11:**
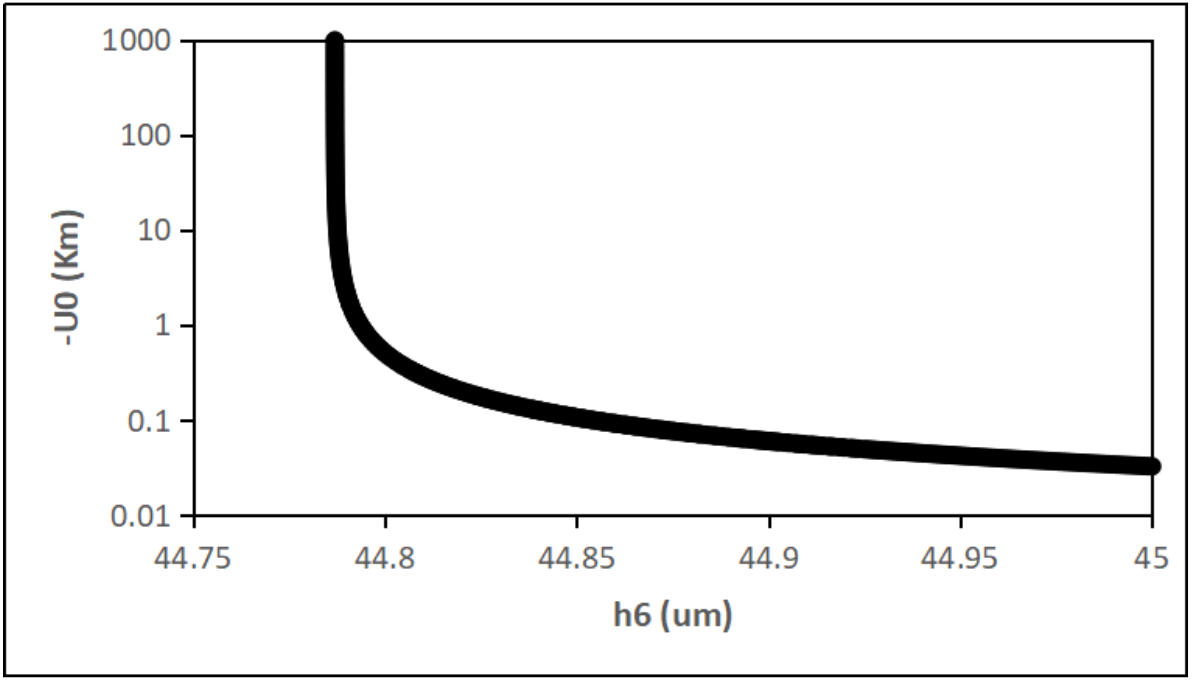
The relationship between the perceivable object distance, *U*_0_, and the interlayer spacing, *h*_6_, between ganglion cell lens array 6 and the bipolar cell lens array, during the perception process from finite to infinite object distances, using a conjugate cone-cell micro-eyepiece array combination 6 (the *U*_0_ axis is expressed in kilometers, log scale).

**Figure 12:**
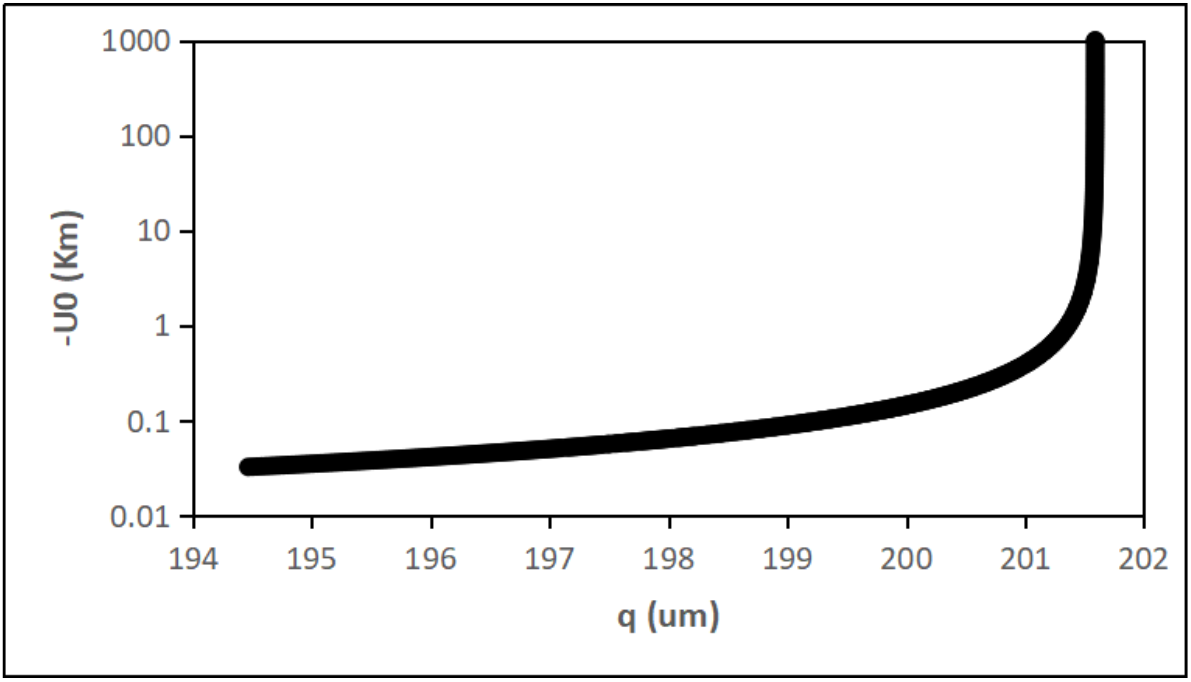
The relationship between the perceivable object distance, *U*_0_, and the distance, *q*, from the bipolar cell lens array of the conjugate mirror to the first photosensitive layer during the perception process from finite to infinite object distances, using a conjugate cone-cell micro-eyepiece array combination 6 (the *U*_0_ axis is expressed in kilometers, log scale).

Table 4,as well as Figures 11 and 12,indicate that a compression of *h*_6_ from 45 to 44.786 µm (214 nm) or a shift of *q* from 194.464 to 201.589 µm (7.125 µm) can enable the cone-cell micro-eyepiece to perceive object distances from a finite distance of 32 m to infinite distances.

Figures 13 and 14 illustrate the relationships between *U*_0_, *h*_6_, and *q* during the process of perceiving object distances from finite to infinite values, calculated using Equations (7) and (8), respectively, when using conjugate rod-cell micro-eyepiece array combination 6.

**Figure 13:**
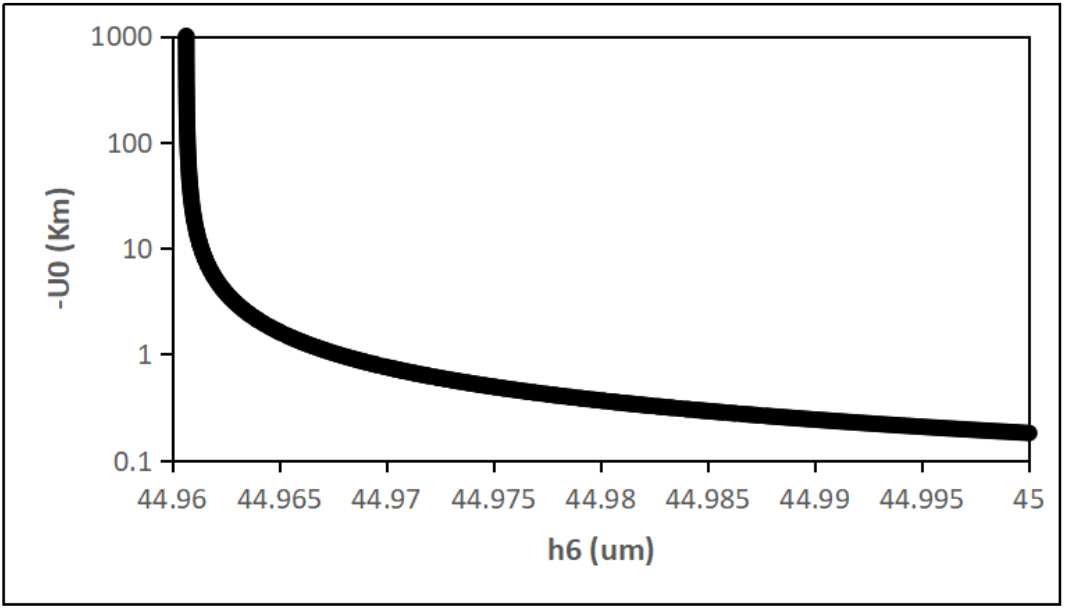
The relationship between the perceivable object distance, *U*_0_, and the interlayer spacing, *h*_6_, between the ganglion cell lens array (layer 6) and the bipolar cell lens array during the perception process from finite to infinite object distances, using a conjugate rod-cell micro-eyepiece array combination 6 (the *U*_0_axis is expressed in kilometers, log scale).

**Figure 14:**
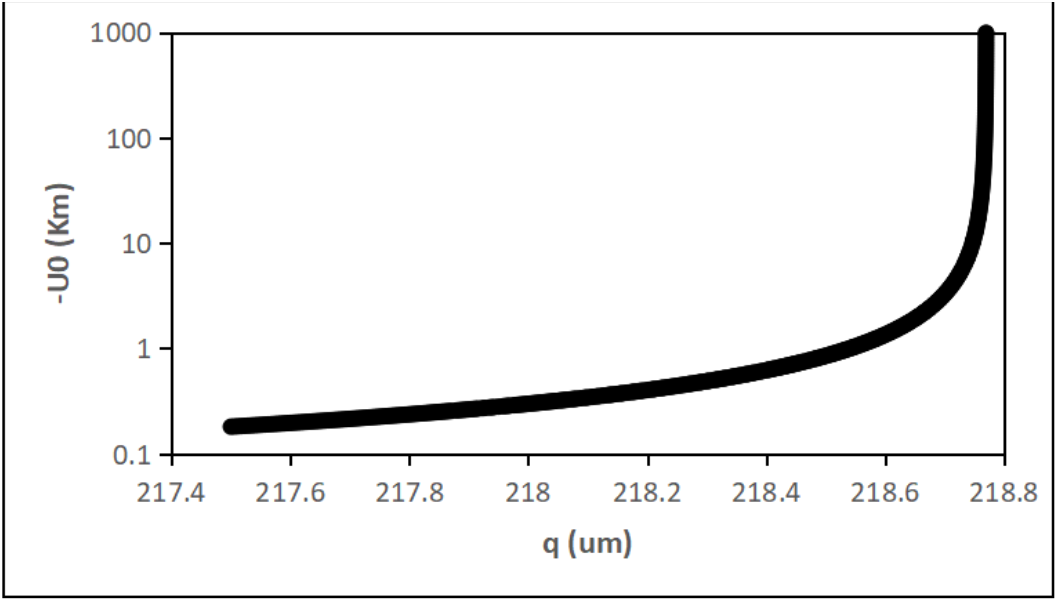
The relationship between the perceivable object distance, *U*_0_, and the distance, *q*, from the bipolar cell lens array of the conjugate mirror to the first photosensitive layer during the perception process from finite to infinite object distances, using a conjugate rod-cell micro-eyepiece array combination 6 (the *U*_0_ axis is expressed in km, log scale).

Table 4,Figures 13 and 14 illustrate that the compression of *h*6 from 45 to 44.960 µm (40 nm), or the displacement of *q* from 217.5 to 218.769 µm (1.269 µm), can enable the perceivable object distance of the rod-cell micro-eyepiece to shift from a finite distance of 180 m to infinite distances.

## 3. Discussion

### 3.1. On the Plausibility of the Assumption that Cell Nuclei Act as Microlenses and that Arrays of Cell Nuclei Form Conjugate Mirrors

The fundamental assumptions of this study is that the retinal ganglion cell nuclei, bipolar cell nuclei, and photoreceptor cell nuclei (cone and rod cell nuclei) also function as optical elements, specifically as optical microlenses or cellular microlenses. The validity of this assumption is directly impacts the feasibility of the three-dimensional imaging model proposed in this paper.

Regarding the optical functions or properties of cell nuclei, the author has not found specific references in the literature. However, the density of cell nuclei is likely higher than that of their surrounding environment. The spherical shape of the nuclei fulfill with the geometric requirements of a lens. Additionally, the diameter of cell nuclei is larger than the wavelength of visible light, making geometric optics theory applicable. Therefore, the assumption is of cellular microlenses appears to be plausible.

The geometric dimensions, optical property values, distribution of various cellular lenses on the retina, the number and arrangement of cellular lens arrays, their interlayer spacing, and other parameters as described in this paper, are speculative. However, they are likely close to actual values. Additionally, these values may vary or change among different individuals and with age in the same individual.

The conjugate mirror system composed of two cellular microlens arrays in the three-dimensional imaging model proposed in this paper is a key optical component for three-dimensional real image formation. The discovery of such conjugate mirrors dates back to the 1970s [6]–[9]. Anatomical analyses of the retina [4], [5] also indicate that ganglion cells and bipolar cells exist in pairs. Therefore, the assumption that a pair of microlens arrays formed by the nuclei of ganglion and bipolar cells constitutes a conjugate mirror system is plausible.

### 3.2. Speculations on the Functions of the Fovea and Macula on the Human Retina

The authors of this article have proposed the following hypothesis regarding the function of the fovea and the macula on the human retina. As shown in Figure 15,the diagram illustrates the optical imaging functionality of the conjugate mirrors on both sides of the fovea. Within the fovea, there are no ganglion cells or bipolar cells present [4,10]. Their absence may reduce the scattering of light caused by the nuclei of these types of cells, resulting in a clearer and brighter conjugate image of the first image. Although there are no ganglion cell lenses or bipolar cell lenses within the fovea to form conjugate mirrors, and even an absence conjugate mirrors altogether, as shown in Figure 15,light from the first image going in other directions can still form a conjugate image at the conjugate position through the conjugate mirrors on the two sides. However, since the light forming the conjugate image originates from the conjugate mirrors on the two sides, it does not include light from the direction where the original object point, the center of the objective lens, and the center of the photoreceptor micro-eyepiece lie on a straight line. The authors hypothesize that the nanoparticles constituting the macula may scatter light from the conjugate image, thereby providing light from this direction to reach the photoreceptor micro-eyepieces and micro-object-distance sensors located in the same direction.

**Figure 15:**
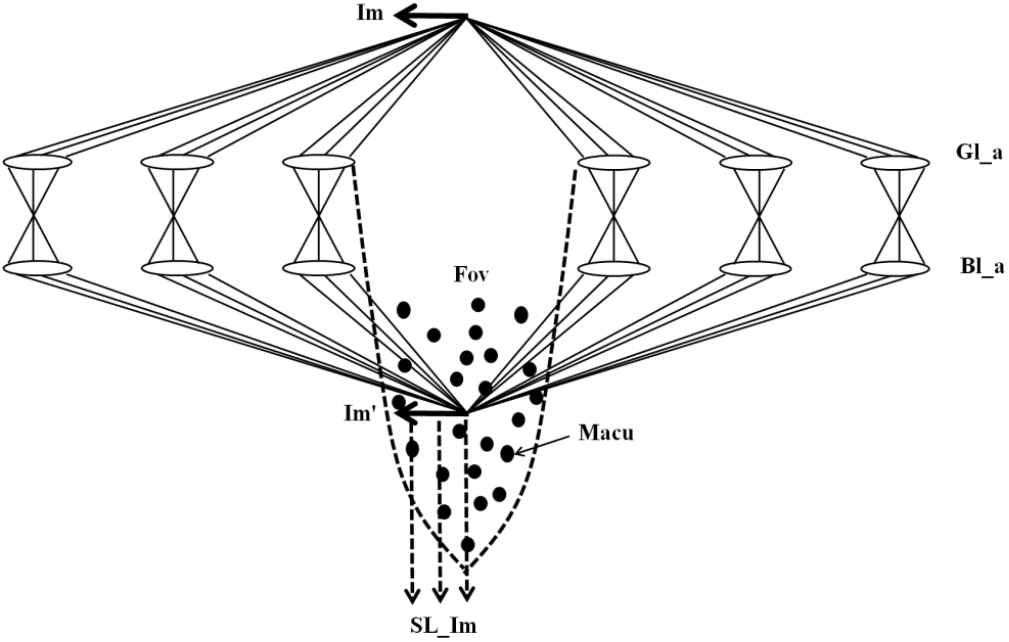
Schematic diagram of the optical imaging functionality of the conjugate mirrors on both sides of the fovea (**Fov**). (The meanings of the symbols in the figure: **Fov**, fovea; **Im**, the first image formed by the objective lens; **Im**’, conjugate image of **Im**; **SL_Im**, scattered light from the conjugate image; **Gl_a**, ganglion cell lens array; **Bl_a**, bipolar cell lens array; **Macu**, nanoparticles of the macula.)

### 3.3. On the Reasonableness of the Hypothesis that the Human Brain Calculates Near to Finite Object Distances by Sensing the Objective Lens Focal Distance When the Photocurrent Disappears

The outer segment of photoreceptor cells, or micro-object-distance sensors, can sense photocurrents, and the magnitude of these photocurrents corresponds to the brightness of the object, as has been extensively reported in the literature [10]. However, in existing research, the photocurrent information generated by the inner segment of photoreceptor cells does not contain depth information across different layers. It is widely accepted that the human eye focuses on objects at varying distances by adjusting the focal length of the crystalline lens [11]. Furthermore, the human brain is capable of sensing and regulating the focal length of the crystalline lens.

This article hypothesizes that real three-dimensional spatial images are formed on photoreceptive membranes at different depths in the micro-object-distance sensors, and these images stimulate the membrane layers at the corresponding depths to generate photocurrents. The depth of the photoreceptive membrane that generates the photocurrent indirectly represents the object distance of the three-dimensional spatial object in the external environment. The human brain first senses the presence of an object in front of the eye by detecting the generation of photocurrents at deeper layers. Subsequently, the brain senses the focal distance of the objective lens at the moment when the photocurrent disappears to calculate the object distance. Therefore, the hypothesis proposed in this article—that the human brain can sense the focal distance associated with the disappearance of photocurrents—is plausible. Additionally, since the brain requires a response time to sense photocurrents, the authors believe that real images first form at deeper layers of the outer segment of photoreceptor cells, giving the brain sufficient time to sense the focal distance at which the photocurrent disappears.

### 3.4. On the Reasonableness of the Assumption That Ganglion Cells and Bipolar Cells Can Move Vertically as a Whole

The computational simulations presented in this article suggest that during the process of perceiving object distances from nearby to finite distances, at the foveal position, the ganglion cell layer and bipolar cell layer—forming the conjugate mirror—can move vertically as a whole relative to the photoreceptor cell layer. The range of this vertical movement is calculated to be 0– 708.5 µm. In contrast, at a position approximately 10 mm away from the fovea, the vertical movement range is much smaller, calculated to be 0–22.5 µm. For positions far from the fovea, if the depth of field of the conjugate imaging by the conjugate mirror is greater than 22.5 µm, the conjugate mirror may still be able to perceive object distances ranging from 20 m to 180 m without requiring vertical movement.

The computational simulations further indicate that during the process of perceiving object distances from finite to infinite distances, at both the foveal position and positions far from the fovea, the interlayer spacing between the ganglion cell layer and the bipolar cell layer—forming the conjugate mirror—needs to undergo slight compression. Specifically, the compression is calculated to be 214 nm in the foveal region and 40 nm in regions far from the fovea.

Additionally, these layers need to move vertically relative to the photoreceptor nuclear layer. The vertical movement is calculated to be 7.125 µm in the foveal region and 1.269 µm in regions far from the fovea. This assumption that the ganglion cell layer and bipolar cell layer can move vertically as a whole, as well as undergo slight interlayer spacing compression, is supported by these computational simulations and provides a plausible explanation for how the human eye might perceive object distances over a wide range.

It is already known that the ganglion cell layer, which constitutes the conjugate mirror, is adjacent to the vitreous body [12]. The vitreous body is soft and should allow the ganglion cell layer to move up and down. Anatomical studies of the retina [4] indicate that the bipolar cell layer and the photoreceptor layer are connected via Henle fibers. At the fovea, this fiber layer is thicker and the fibers are longer compared to the peripheral retina. Anatomical photographs of the fovea show that these Henle fibers lie flat on the retina. Therefore, this paper hypothesizes that the process of the conjugate mirror moving up and down as a whole relative to the micro-eyepiece array/photoreceptor layer may be accomplished by the Henle fibers transitioning from a flat to an upright position.

### 3.5 Other

The following hypotheses and speculations in this paper require further investigation:

1. Can a single micro-real-image telescope or a single photoreceptor perceive the object distance of a single object point in one azimuthal direction (defined by two angular coordinates), or can it perceive the object distances of multiple object points within a small angular range?
2. Does the human brain indeed perceive object distance by detecting the focal length of the objective lens or focal plane spacing (the distance between the conjugate focal plane of the objective lens and the object-side focal plane of the micro-eyepiece) when the photocurrent disappears, or is it through some other process?
3. When the human brain perceives the object distance of a finitely distanced object, it is assumed that during the process of the objective lens changing its focal length, the center of the objective lens does not move, and the total distance between the center of the objective lens and the first photosensitive layer of the micro-object distance sensor remains constant. However, this assumption might be relaxed. As long as the distance between the image-side principal plane of the objective lens and the first photosensitive layer remains constant (this is achieved by moving the center of the objective lens), it should still be possible to perceive the object distance of a finitely distance object. Apart from this hypothesized process, could there be other mechanisms involved?
4. Can the human brain perceive and control the focal plane spacing, including the ability to perceive and control the focal plane spacing to be zero (where the conjugate focal plane of the objective lens coincides with the image-side focal plane of the micro-eyepiece)?
5. Can the human brain perceive and control the distance between the bipolar cells and the photoreceptor cells?
6. Can the human brain perceive and compress the minute distance between the ganglion cells and the bipolar cells?

However, these discussions do not affect the validity of the three-dimensional imaging optical model itself. They only influence the hypotheses about how the human brain uses and controls this three-dimensional imaging optical model to perceive object distance.

The monocular stereoscopic vision model proposed in this paper does not conflict with the well-known binocular stereoscopic vision model. For nearby objects, the image-side azimuthal angles on the retinas of the two eyes, with the center of each eye’s objective lens as the origin, differ. The binocular stereoscopic vision model perceives object distance based on the difference in these image-side azimuthal angles (parallax). The closer the object, the greater the parallax. However, when the object is far enough away, the parallax becomes smaller than the spacing between photoreceptor cells, making it imperceptible to the brain. At this point, the binocular stereoscopic vision model fails. In contrast, the monocular stereoscopic vision model proposed in this paper allows a single eye to perceive objects at distances ranging from 25 cm to infinite distances. The difference in azimuthal angles between the left and right eyes for nearby objects does not affect the ability of a single eye to perceive these objects. Each eye independently perceives object distance. The author of this paper speculates that the combination of binocular stereoscopic perception for near distances and the monocular stereoscopic perception of each eye may work together to improve the accuracy of distance perception for nearby objects. At close distances, binocular vision can perceive a wider range of object-side azimuthal angles compared to monocular vision.

Additionally, this article hypothesizes that the human eye, even with just one eye (monocular vision), can achieve stereoscopic vision. Combined with the ability of a single eye to perceive colored objects, it is thus proposed that the human eye, when using one eye alone, possesses colored stereoscopic vision.

In the three-dimensional imaging model presented in this article, the image on the outer segment of the photoreceptor cells is upright. The commonly accepted notion of an inverted image on the retina, which corresponds to an intermediate image in our model. All assumptions about the three-dimensional real-image imaging model and the estimates used in the computational simulations presented in this article differ from actual values to some extent, and their validity remains to be verified by scientists in the relevant fields.

The micro-real-image telescope proposed in this article can be compared to the conventional virtual-image telescope when the distance *D* between the conjugate focal plane of the objective lens and the focal plane of the micro-eyepiece vanishes. The difference lies in the nature of the image: one forms a real image, while the other forms a virtual image. Assuming the maximum focal length of the objective lens (the human cornea + lens) is 18.4 mm (with an image-side focal length of 24.5 mm), and the focal lengths of the micro-eyepieces formed by cone cell nuclei and rod cell nuclei are 20 μm and 10 μm, respectively, we can use the equation 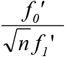 for the magnification of a telescope. We then find that a cone-cell micro-real-image telescope can achieve a magnification of up to 1062x, while a rod-cell micro-real-image telescope can achieve 2124x. These are quite high magnifications for telescopes.

### 3.6. Possible Applications

If the three-dimensional imaging optical model and stereoscopic perception process proposed in this article are validated, they could contribute to the following areas:

1. Treatment of Eye Diseases and Blindness The model could provide valuable insights for the treatment of vision-related eye diseases and blindness. It might even inspire the design and creation of artificial eyes, similar to artificial cochleae, to replace damaged eyeballs and retinas. These artificial eyes could generate electrical signals in a format that is compatible with biological optical nerves.
2. Development of Three-Dimensional Cameras for Human Tissue and Organs The model could inspire the design and development of three-dimensional cameras capable of imaging human tissues, organs, and blood vessels. Such cameras could utilize infrared light sources that are safe for the human body and can partially penetrate human tissues [13].
3. Cancer Treatment Devices The model could inspire the development of equipment for cancer treatment. Considering the reciprocity of optical imaging—where the positions of the light source and the imaging location can be interchanged—humans could design medical devices that combine three-dimensional imaging and light-concentration therapy. For example, such a device could emit laser beams at the three-dimensional image location of cancerous tumor blood vessels. The laser would focus at the physical location of the blood vessels, cauterizing and blocking blood flow, thereby cutting off the energy supply to the cancerous tumor. This could lead to the death of cancer cells, achieving the goal of cancer treatment.
4. Inspiration for Three-Dimensional Range-Finding Cameras The model could inspire the design of three-dimensional range-finding cameras, which could be used to develop stereoscopic vision systems for autonomous vehicles. These systems would mimic the way the human eye perceives the world in three dimensions.
5. If we can design eyepieces for astronomical telescopes with focal lengths in the tens of µm range, arranged as micro-eyepiece arrays, we could potentially increase the magnification of existing astronomical telescopes by nearly a thousand times, while also achieving an omnidirectional 360° field of view.

## Conclusion

This article proposes a geometric optical model for three-dimensional real-image formation in monocular vision, based on the assumption that the ganglion cell nuclei, bipolar cell nuclei, and photoreceptor cell nuclei (cone and rod cell nuclei) in the human retina function as optical microlenses and actively participate in optical imaging.

The model assumes that our monocular vision operates as an array of high-magnification micro-real-image telescopes oriented toward numerous azimuthal directions (each azimuth defined by two angular coordinates). These micro-real-image telescopes are composed of the following components:

- The cornea and crystalline lens, which serve as objective lenses with tunable focal lengths,
- Paired arrays of ganglion cell nuclei and bipolar cell nuclei, which act as conjugate mirrors,
- Individual photoreceptor cell nuclei (cone or rod nuclei), which function as micro-eyepieces,
- The outer segment of each photoreceptor cell, which serves as a micro-object-distance sensor. The two angular coordinates of a single micro-eyepiece correspond to the two angular coordinates of the azimuth of a three-dimensional object in space. The photoreceptor outer segment, as the sensor for each micro-real-image telescope, consists of multiple photosensitive layers at varying depths, with the depth corresponding to object distance in three-dimensional space.

This text hypothesizes that the human eye comprises six combinations of conjugate mirrors and micro-eyepiece arrays. Each conjugate micro-eyepiece combination is responsible for perceiving object distances within different ranges, from near to finite distances. The sixth combination is capable of perceiving object distances from the farthest finite distances to nearly infinite distances.

The text further hypothesizes that the process by which one human eye perceives object distances involves two stages: from near to finite distances, and from finite distances to infinite distances. In both stages, the brain maintains the distance between the optical center of the cornea and crystalline lens (treated as the objective lens) and the center of the micro-eyepiece array. In the process of perceiving object distances from near to finite distances, the brain ensures that the conjugate focal plane of the objective lens coincides with the object-side focal plane of the micro-eyepiece array (that is, the focal plane spacing is zero). Perception is then achieved by observing the process during which the focal length of the objective lens increases. Specifically, perception occurs when the photocurrent generated by the optical system, starting from a nonzero value, diminishes, and the focal length of the objective lens at the moment of photocurrent disappearance determines the object distance. In the process of perceiving object distances from finite distances to infinite distances, the brain maintains the focal length of the objective lens at the maximum focal length of the sixth conjugate micro-eyepiece combination, where the focal plane spacing is zero. Perception is then achieved by observing the process during which the focal plane spacing increases from zero. Specifically, perception occurs when the photocurrent transitions from being present to absent (or becomes faintly perceptible), and the focal plane spacing at the moment of photocurrent disappearance (or faint appearance) determines the object distance. When the focal plane spacing reaches a fixed value, the perceived object distance is infinite. However, the brain’s ability to distinguish differences in focal plane spacing determines the maximum resolvable object distance.

The computational simulation of object distance perception described in this text suggests that the nucleus of a cone cell around the fovea acts as a micro-eyepiece in a micro-real-image telescope. By scanning the focal length of the objective lens from 15.9 mm to 24.7 mm, the human brain can perceive object distances ranging from a nearest distance of 25 cm to a maximum distance of 32.12 m. Similarly, the nucleus of a rod cell in the peripheral retina acts as a micro-eyepiece in a micro-real-image telescope. By scanning the focal length of the objective lens from 16.0 mm to 24.6 mm, the human brain can perceive object distances ranging from a nearest distance of 20.2 m to a maximum distance of 180.58 m. When the conjugate focal plane of the objective lens is separated from the focal plane of the cone and rod-cell micro-eyepiece by distances of 14.035 µm and 2.500 µm respectively, it becomes possible to perceive objects at distances ranging from 32.12 m and 180.58 m to infinity. If the human brain is capable of distinguishing differences in focal plane spacing as small as 1 nm, 10 nm, or 20 nm, the micro-real-image telescope can differentiate objects at distances of 450 km, 45 km, or 23 km, respectively, from objects at infinite distances.

## Acknowledgments

The authors of this article would like to express their gratitude to Ms. Lü Pinjing of Jmuse Technologies Co., Ltd. for her invaluable assistance! Ms. Lü meticulously created Figure 1,Figure 1A,and Figure 1B for this article.

